# Characterization of CAP1 and ECA4 adaptors participating in clathrin-mediated endocytosis

**DOI:** 10.1101/2022.01.07.475412

**Authors:** Maciek Adamowski, Ivana Matijević, Jiří Friml

## Abstract

Formation of endomembrane vesicles is crucial in all eukaryotic cells and relies on vesicle coats such as clathrin. Clathrin-coated vesicles form at the plasma membrane and the *trans-*Golgi Network. They contain adaptor proteins, which serve as binding bridges between clathrin, vesicle membranes, and cargoes. A large family of monomeric ANTH/ENTH/VHS adaptors is present in *A. thaliana*. Here, we characterize two homologous ANTH-type clathrin adaptors, CAP1 and ECA4, in clathrin-mediated endocytosis (CME). CAP1 and ECA4 are recruited to sites at the PM identified as clathrin-coated pits (CCPs), where they occasionally exhibit early bursts of high recruitment. Subcellular binding preferences of N- and C-terminal fluorescent protein fusions of CAP1 identified a functional adaptin-binding motif in the unstructured tails of CAP1 and ECA4. In turn, no function can be ascribed to a double serine phosphorylation site conserved in these proteins. Double knockout mutants do not exhibit deficiencies in general development or CME, but a contribution of CAP1 and ECA4 to these processes is revealed in crosses into sensitized endocytic mutant backgrounds. Overall, our study documents a contribution of CAP1 and ECA4 to CME in *A. thaliana* and opens questions about functional redundancy among non-homologous vesicle coat components.

## Introduction

In plants, like in other eukaryotes, the formation of vesicles at the internal compartments of the endomembrane system, as well as at the plasma membrane (PM), depends on the activity of a plethora of proteins recruited from the cytosol and together constituting vesicle coats (Béthune and Wieland, 2018; Kaksonen and Roux, 2018). Clathrin-coated vesicles (CCVs), generated at the PM as well as at the *trans*-Golgi Network (TGN), employ the coat protein clathrin, which forms a cage-like structure around the nascent vesicle (Fotin et al., 2004; Narasimhan et al., 2020). The clathrin coat polymerizes from building blocks called triskelia (sing. triskelion), heterohexamers composed of three CLATHRIN LIGHT CHAIN (CLC) and three CLATHRIN HEAVY CHAIN (CHC) subunits.

Besides clathrin, protein coats around forming CCVs prominently contain so-called adaptor proteins (Owen et al., 2004). This class of proteins is named so due to their function as mediators of the binding between the membrane of the nascent vesicle and/or cargoes packed into the vesicle, and between clathrin, which does not directly interact with membranes. Adaptor proteins share a common design principle, namely, are composed of one or more folded domains and of unstructured regions. The folded domains interact with lipids of the membranes, and contain binding pockets for short peptide motifs present in cargoes and in other coat components. The unstructured regions serve to accommodate the protein within the coat structure, and contain short peptide motifs binding with clathrin and with the folded domains of other coat components (Owen et al., 2004). Despite this common design, reflecting the role of adaptors within vesicle coats, adaptor proteins belong to several phylogenetically distinct classes. Beside the relatively well-characterized heterotetrameric adaptor protein complexes (APs), of which AP-1, AP-2, and AP-3 are known to associate with clathrin (Park and Guo, 2014; Sanger et al., 2019), a major adaptor class found both in plants and in other eukaryotes is the ANTH/ENTH/VHS superfamily (Owen et al., 2004; Itoh and De Camilli, 2006; Zouhar and Sauer, 2014). These are monomers composed of a single folded ANTH (AP180 N-Terminal Homology), ENTH (Epsin N-Terminal Homology), or VHS (Vps-27, Hrs, and STAM) domain at the N-terminus, followed by an unstructured region of a varied length and composition. The major mammalian ANTH isoforms are AP180 and CALM (clathrin assembly lymphoid myeloid leukemia protein), while major ENTH adaptors are called epsins (Owen et al., 2004). The ANTH and ENTH domains are distinct, but form similar folds, and both bind to phosphatidylinositol 4,5-bisphosphate (Itoh et al., 2001; Ford et al., 2002). The function of ANTH/ENTH/VHS adaptors at clathrin-coated pits (CCPs) includes a contribution to membrane curvature, which may be effected either by the insertion of amphipathic helices, as classically described in epsins (Ford et al., 2002; Stahelin et al., 2003) and more recently in CALM (Miller et al., 2015), or alternatively, by protein crowding, which produces steric pressure that drives membrane bending, especially in the case of disordered proteins (Stachowiak et al., 2012, 2013; Busch et al., 2015). *In vitro,* ANTH and ENTH proteins form co-assemblies, which may also be a feature of their common organization in genuine CCPs *in vivo* (Skruzny et al., 2015; Garcia-Alai et al., 2018). Finally, both these adaptor types contribute to the fusion of the complete vesicle with its target by recruiting the vesicle fusion proteins SNAREs (SNAP receptors) (Miller et al., 2007, 2011).

In plants, the ANTH/ENTH/VHS superfamily is expanded, with 35 predicted proteins in *Arabidopsis thaliana* (Zouhar and Sauer, 2014). Characterization of family members reveals diverse roles in endocytic, secretory, and vacuolar trafficking. EPSIN1 and MTV1 (Song et al., 2006; Sauer et al., 2013), TGN-localized ENTH adaptors in *A. thaliana*, lack close homology and localize to distinct subdomains of the organelle, where they preferentially associate with CCVs containing the adaptor complexes AP-1 and AP-4, respectively (Heinze et al., 2020). Despite these molecular differences, EPSIN1 and MTV1 are functionally redundant in vacuolar trafficking, and contribute to secretory traffic as well (Heinze et al., 2020). Other ENTH homologues, EPSIN2 and EPSIN3, localize to the PM (Heinze et al., 2020), although EPSIN2 has been previously reported to associate both with AP-2, which participates in CME, and with AP-3, which acts in vacuolar trafficking (Lee et al., 2007; Zwiewka et al., 2011; Di Rubbio et al., 2013).

Within the ANTH subfamily of *A. thaliana*, PICALM5a/ECA2 and PICALM5b, as well as PICALM9b/EAP1, are involved in the function of pollen tubes (Muro et al., 2018; Li et al., 2018), where the homologous PICALM5 pair controls the pollen tube tip localization of ANXUR receptor kinases required for membrane integrity (Muro et al., 2018). In turn, PICALM1a/ECA1 and PICALM1b function in the PM retrieval of VAMP72 SNAREs, proteins required for secretory vesicle fusion (Fujimoto et al., 2020). Finally, the homologous pair PICALM4b/CAP1 (CAP1 in the following) and PICALM4a/ECA4/AtECA4 (ECA4 in the following) was found to physically interact with the endocytic TPLATE complex, which presumably acts as a clathrin adaptor itself (Gadeyne et al., 2014). CAP1 was also identified as an interactor of CLC and of AUXILIN-LIKE1/2, putative uncoating factors for endocytic CCVs (Adamowski et al., 2018). Both CAP1 and ECA4 localize to the PM and intracellular compartments, although CAP1 is found at the latter only rarely (Song et al., 2012; Nguyen et al., 2018; Adamowski et al., 2018). A role of ECA4 in cargo recycling from the TGN to the PM was suggested by the study of *eca4* mutants (Nguyen et al., 2018), but overall, the role of CAP1 and ECA4 in endomembrane trafficking, and in particular in CME, is not well understood.

Here, to extend the understanding of the ANTH adaptors CAP1 and ECA4, we characterize these proteins on the molecular and functional levels with a focus on a potential role in CME. We analyze the CAP1 protein composition, visualize CAP1 and ECA4 localization at the PM with Total Internal Reflection Fluorescence (TIRF) microscopy, and characterize *cap1 eca4* mutants, also in crosses with mutants of other components of CME.

## Results

### Recruitment of CAP1 and ECA4 to CCPs at the PM

CAP1 and ECA4 have been previously localized to the PM and to intracellular structures (Song et al., 2012; Nguyen et al., 2018; Adamowski et al., 2018). The intracellular site of ECA4 localization has been identified as the TGN (Nguyen et al., 2018). CAP1 has been detected at intracellular structures variably, while the PM appears to be its predominant site of action (Adamowski et al., 2018). To characterize the activity of CAP1 and ECA4 at the PM in more detail, we employed TIRF microscopy in a cross between *CsVMV_pro_:ECA4-GFP* and *CAP1_pro_:CAP1- mCherry* fluorescent protein markers. With TIRF time lapses in the epidermis of etiolated hypocotyls, we observed both CAP1-mCherry and ECA4-GFP recruitment into distinct structures at the PM surface, where they persisted for varied periods of time (Figure 1A). CAP1-mCherry and ECA4-GFP were often, but not exclusively, co- localized on the same structures. In a cross between a clathrin marker *CLC2_pro_:CLC2-GFP* and *CAP1_pro_:CAP1-mCherry*, we observed that CAP1-mCherry is recruited to forming CCPs, as the signals of the two marker proteins often overlapped in kymographs (Figure 1B). Interestingly, CAP1-mCherry relatively often exhibited a burst of intensive recruitment at the onset of CCP formation (Figure 1B, arrowheads). The level of CAP1-mCherry signal was then relatively low during the subsequent stages up until the moment of vesicle scission from the PM, when the CLC2-GFP trace on kymographs ended. This observation suggests that CAP1 might be particularly active at the early stages of CCP formation. However, in contrast to the relatively frequent detection of such recruitment patterns in the *CLC2_pro_:CLC2- GFP CAP1_pro_:CAP1-mCherry* marker cross, CAP1-mCherry exhibited these early bursts of recruitment only very rarely when expressed alone in *cap1-1 eca4-2* double mutant background (Figure 1C; the double mutant is described below). With an independent *35S_pro_:CAP1-RFP* construct, expressed in the wild type background, we observed this recruitment pattern equally rarely (Figure 1D). The vast majority of PM recruitment events recorded in *cap1-1 eca4-2 CAP1_pro_:CAP1-mCherry* and in *35S_pro_:CAP1-RFP* were characterized by an equal amount of fluorescent signal along the time course of recruitment at distinct foci on the PM (Figure 1C and 1D). ECA4-GFP, expressed in the double marker cross with CAP1-mCherry, too, was found to exhibit initial bursts of high recruitment only in rare cases (Figure 1E).

**Figure 1.**
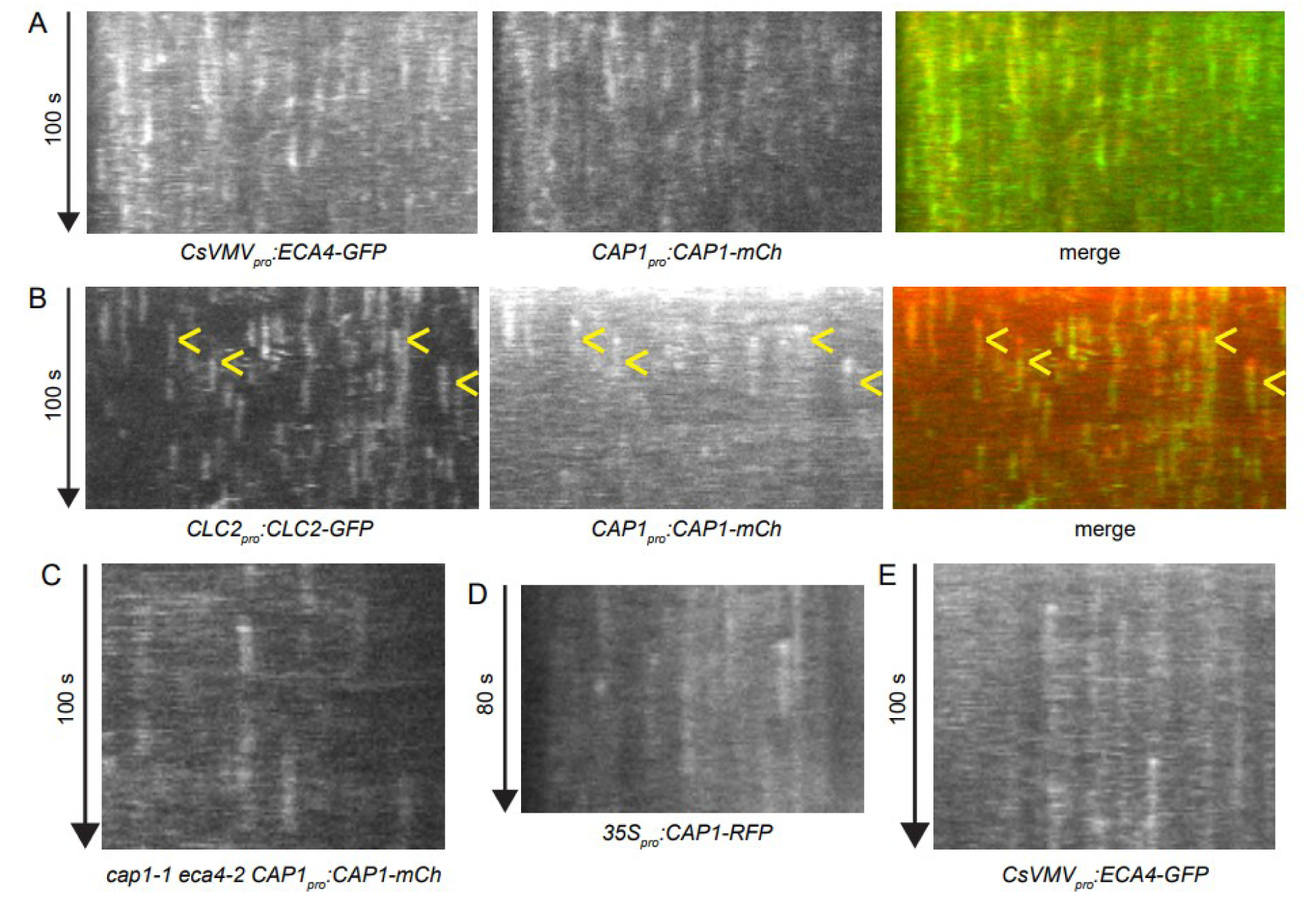
Dynamics of CAP1 and ECA4 recruitment to CCPs. All panels show kymographs of TIRF time lapses captured in the epidermis of etiolated hypocotyls. (A) Co-localization of CAP1-mCherry and ECA4-GFP. CAP1 and ECA4 persist at the PM and often co-localize on the same structures. (B) Co-localization of CAP1-mCherry and CLC2-GFP. CAP1 localizes to CCPs and variably exhibits bursts of recruitment at the initiation of CCP formation. (C, D) Localization of CAP1-mCherry expressed in *cap1-1 eca4-2* mutant background (C) and of CAP1-RFP expressed in wild type background (D). In these lines, CAP1 is mostly observed as traces of equally intense signal, while cases of early bursts of recruitment are rare. (E) ECA4-GFP rarely exhibits early bursts of recruitment, and in most cases exhibits equal recruitment throughout its life time at the PM.

Altogether, TIRF imaging of CAP1 and ECA4 fluorescent reporter fusions establishes that they are recruited to forming CCPs at the PM, and there are indications that the adaptors may occasionally undergo an early burst of recruitment at the initiation of CCP formation. Yet, it is unlikely that this represents their typical mode of action. It is not clear why this recruitment pattern could be seen more readily in a marker cross between *CAP1_pro_:CAP1-mCherry* and *CLC2_pro_:CLC2-GFP*.

### Increased CAP1 recruitment to the PM following overexpression of AUXILIN-LIKE1

Overexpression of AUXILIN-LIKE1/2, putative uncoating factors for CCVs in *A. thaliana*, leads to an inhibition of CME (Adamowski et al., 2018, 2021). The inhibition of CME by AUXILIN-LIKE1/2 overexpression manifests by specific changes in the PM binding patterns of fluorescent reporter fusions of proteins active in the endocytic process. Namely, CLC2-GFP and, to some degree, the dynamin marker DRP1C- GFP (DYNAMIN RELATED PROTEIN 1C; Konopka et al., 2008) become depleted from the PM, while TPLATE-GFP, a marker for a subunit of the TPLATE adaptor complex (Gadeyne et al., 2014), and AP2A1-TagRFP, a marker for a subunit of the AP-2 adaptor complex (Di Rubbio et al., 2013), bind to the PM excessively when AUXILIN-LIKE1 is overexpressed (Adamowski et al., 2018). These observations indicate that overexpression of AUXILIN-LIKE1 inhibits CME at the stage of clathrin recruitment, but after the initial steps, which incorporate the early-arriving TPLATE and AP-2 complexes (Gadeyne et al., 2014) are still functional.

We tested how CAP1-mCherry reacts to the overexpression of AUXILIN- LIKE1 by crossing *CAP1_pro_:CAP1-mCherry* with *XVE»AUXILIN-LIKE1,* a line overexpressing this putative uncoating factor in an inducible fashion upon chemical induction by β-estradiol. Approximately 24 h after expression induction, overexpression of AUXILIN-LIKE1 lead to a strong increase in the PM association of CAP1-mCherry, observed with Confocal Laser Scanning Microscopy (CLSM) in seedling root apical meristem (RAM) epidermis (Figure 2A, 2B). This observation is similar to the increased recruitment of TPLATE-GFP and AP2A1-TagRFP to the PM in analogical conditions (Adamowski et al., 2018), and as such, supports the notion that CAP1 is an early-arriving adaptor protein during CCP formation.

**Figure 2.**
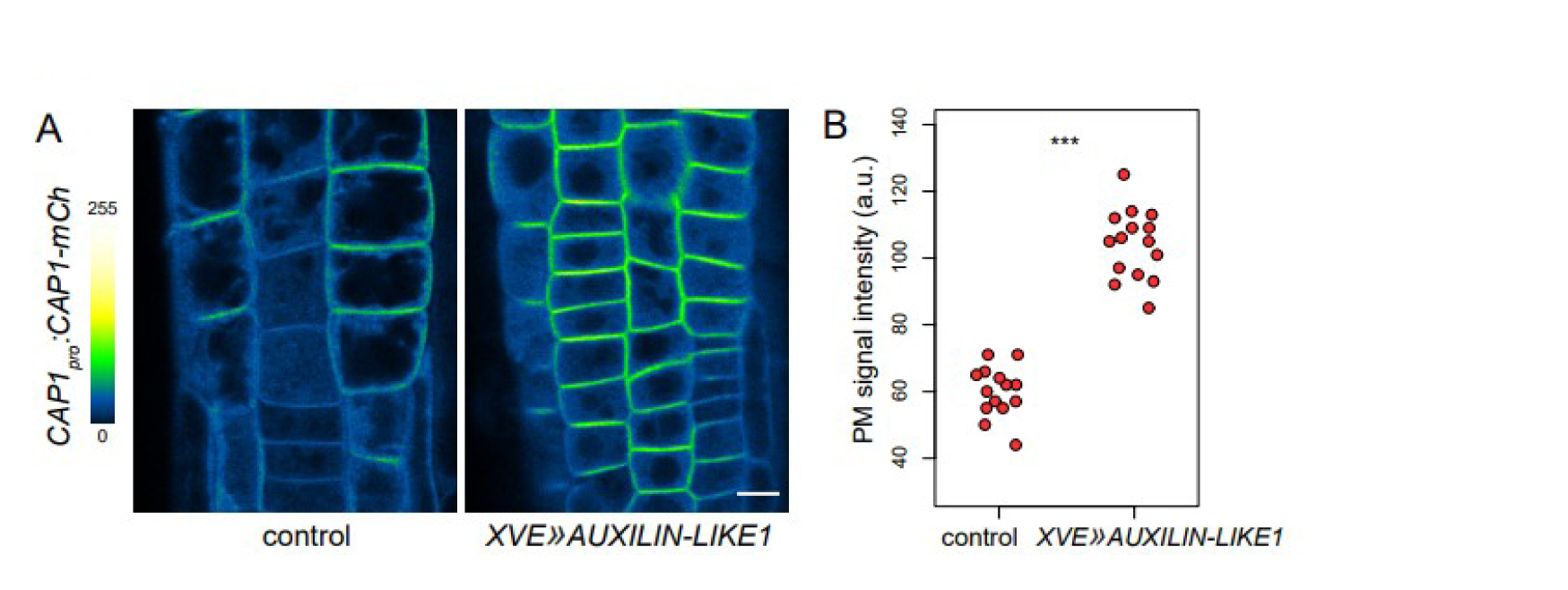
Increased PM binding of CAP1-mCherry following AUXILIN-LIKE1 overexpression. (A) CLSM images of CAP1-mCherry in seedling RAM epidermis in control conditions and after induction of AUXILIN-LIKE1 overexpression for 24 h. Scale bar – 10 µm. (B) Quantification of PM signal intensities of CAP1-mCherry in control conditions and after induction of AUXILIN-LIKE1 overexpression for 24 h from a representative experiment. Each dot represents a measurement from one root. Control: 59.9 ± 7.6 a.u., n=14; *XVE»AUXILIN-LIKE1*: 104 ± 10.4 a.u., n=15. Values were compared using a *t* test, *** – P<0.0001.

### Analysis of CAP1 domains and a putative AP-binding motif within the unstructured region

Both ENTH and ANTH adaptors are composed of two major domains: an N-terminal folded ENTH/ANTH domain, which binds to membranes, and a C-terminal unstructured region of varied composition. The unstructured region often contains short peptide motifs involved in weak molecular interactions with clathrin and with the AP-2 complex (Itoh and De Camilli, 2006; Smith et al., 2017). CAP1 and ECA4 share a very high sequence homology, both in the ANTH domain and in the unstructured region (Suppl. Figure 1). The unstructured regions contain a single FS motif (Phe-Ser; ^452^FS and ^472^FS in CAP1 and ECA4, respectively), identified as a minor clathrin binding motif in the CCV uncoating factor auxilin (Scheele et al., 2003), as well as a single DPF motif (Asp-Pro-Phe; ^555^DPF and ^574^DPF in CAP1 and ECA4, respectively), known for an affinity with a binding pocket in the appendage domain of the α2 subunit of AP-2 (Smith et al., 2017).

**Suppl. Figure 1.**
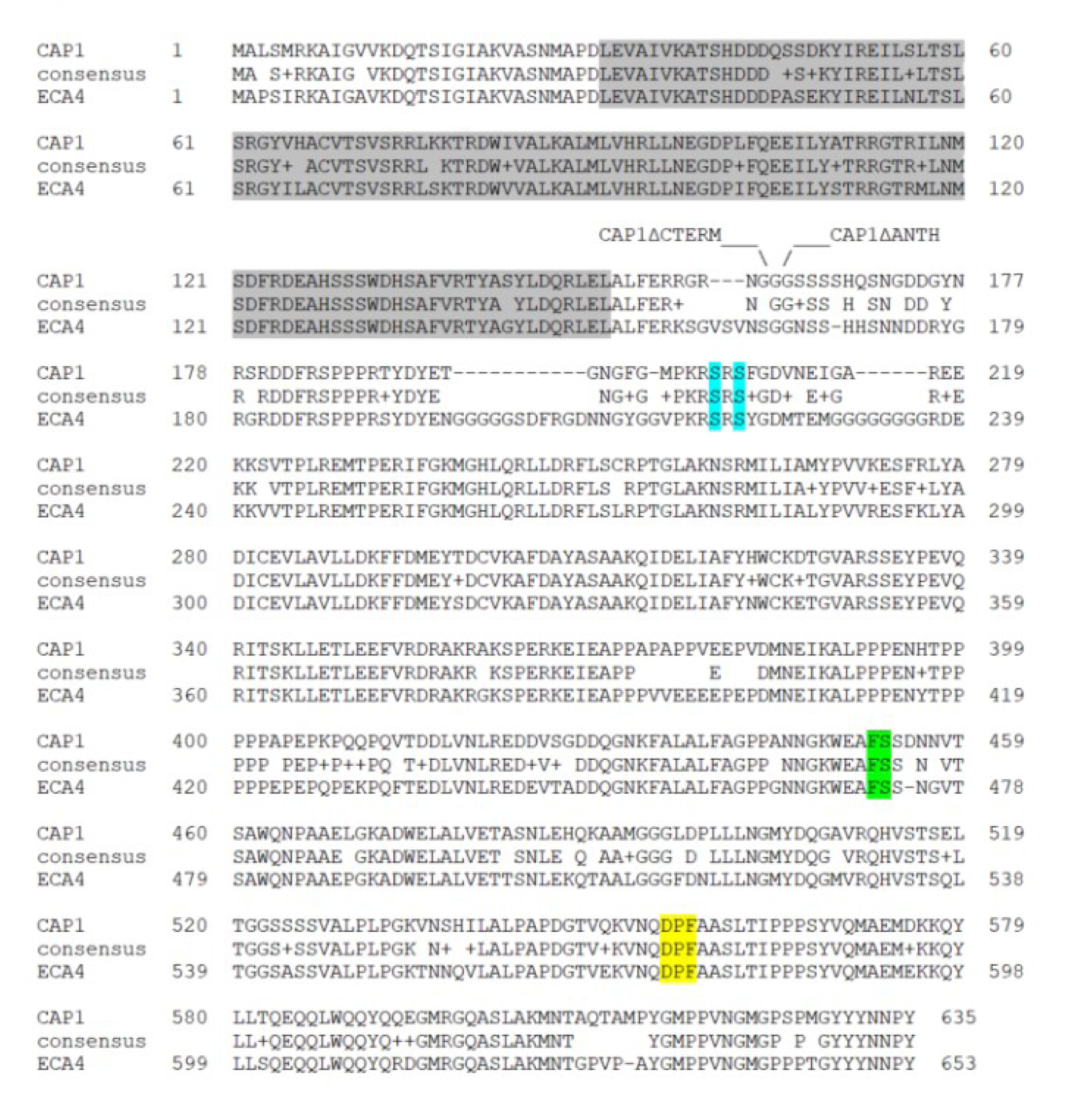
Alignment of CAP1 and ECA4 protein sequences BLAST alignment of CAP1 and ECA4 protein sequences. Sequence encoding the ANTH domain is labelled in grey. Borders of CAP1^ΔCTERM^ and CAP1^ΔANTH^ truncations are indicated. GFP-CAP1^ΔCTERM^ contains amino acids 1-161 and CAP1^ΔANTH^ contains amino acids 163-473. The double serine phosphorylation site is labelled in blue. The FS motif is labelled in green. The DPF motif is labelled in yellow.

To test the molecular functions of ANTH and C-terminal domains of CAP1 and ECA4, we cloned truncated variants of CAP1 and expressed them as fluorescent protein fusions in stably transformed *A. thaliana* to observe subcellular localizations. CAP1^ΔANTH^, which contains only the unstructured C-terminal region, and CAP1^ΔCTERM^, which contains only the ANTH domain, were expressed as fusions with mCherry at the C-terminus, in constructs analogical to the full length *CAP1_pro_:CAP1- mCherry*. In contrast with full length CAP1-mCherry, localizing typically to the PM (Figure 3A), CAP1^ΔCTERM^-mCherry was bound to intracellular compartments, or, possibly, was present as intracellular protein aggregates, which did not colocalize with TGN-localized clathrin labeled by CLC2-GFP (Figure 3B). In turn, CAP1^ΔANTH^- mCherry did not express in any of the transgenic lines tested. These results indicate the requirement of both major domains of the CAP1 protein for its correct expression and subcellular localization.

**Figure 3.**
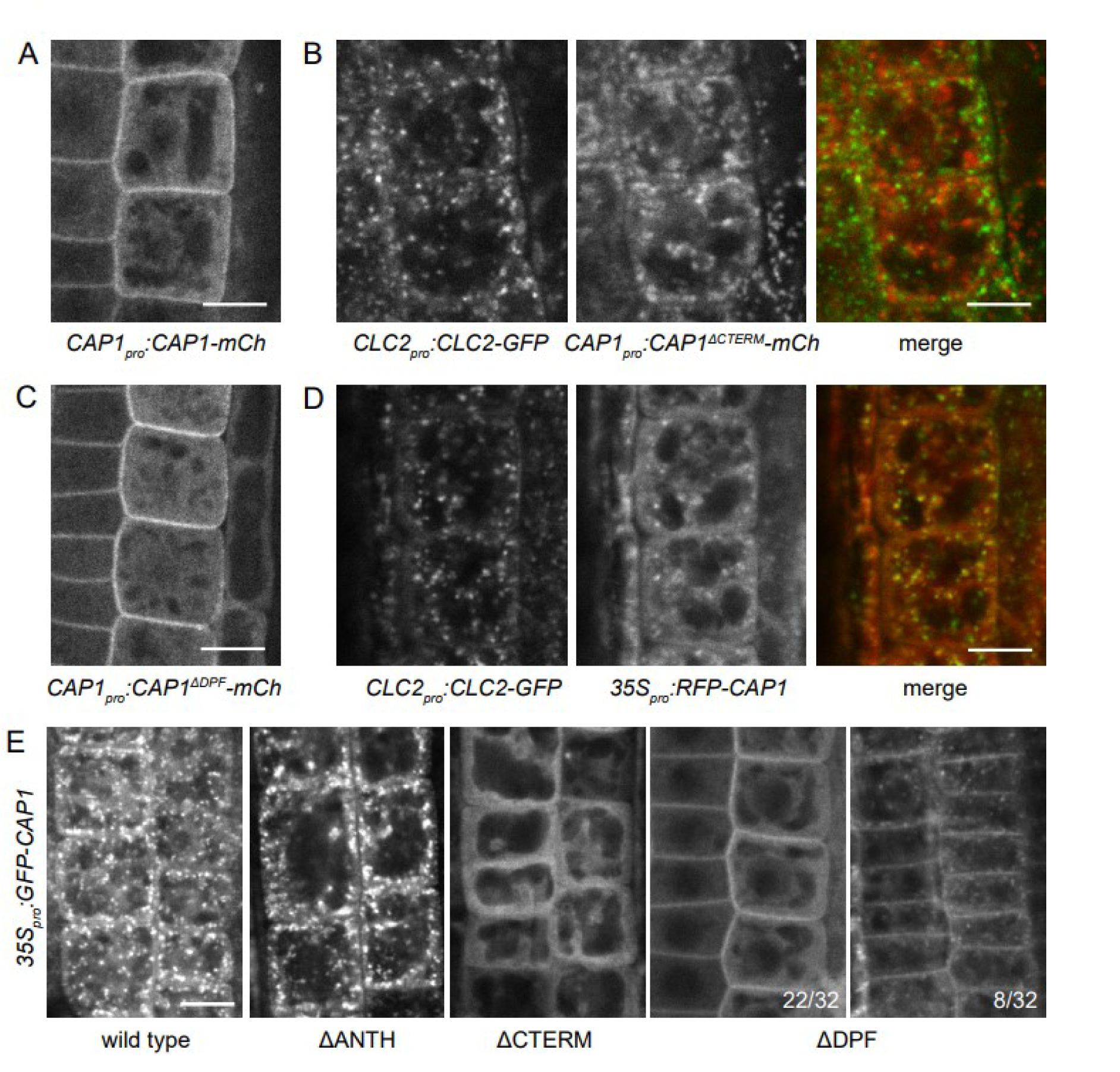
Subcellular localizations of CAP1 fluorescent protein fusions and deletion constructs. All panels show CLSM images taken in seedling RAM epidermis. Scale bars – 10 µm. (A) CAP1-mCherry typically localizes to the PM. (B) CAP1^ΔCTERM^-mCherry localizes to intracellular structures or protein aggregates distinct from TGNs labelled by CLC2-GFP. (C) CAP1^ΔDPF^-mCherry typically localizes to the PM. (D) RFP-CAP1 localizes to TGNs labelled by CLC2-GFP. (E) Comparison of wild type GFP-CAP1 and deletion constructs. GFP-CAP1 and GFP-CAP1^ΔANTH^ localize to intracellular structures, presumably TGN. GFP-CAP1^ΔCTERM^ does not bind to any membranes. GFP-CAP1^ΔDPF^ localizes to the PM, in some cases accompanied by localization to intracellular structures. Numbers indicate ratios of RAMs with particular localization patterns of GFP- CAP1^ΔDPF^. In 2 samples, GFP-CAP1^ΔDPF^ localized only to intracellular structures (not shown).

Similarly, we tested the molecular function of the DPF motif by generating a mutant variant CAP1^ΔDPF^ where ^555^DPF is substituted with alanines for ^555^AAA. CAP1^ΔDPF^-mCherry exhibited a normal localization at the PM (Figure 3C), not indicative of a function of the ^555^DPF sequence in the recruitment of CAP1 to CCPs through possible interactions with AP-2.

Additionally, we generated fusions of wild type CAP1 and the above-described mutant variants, with fluorescent proteins attached at the N-terminus (collectively designated FP-CAP1). Fusions of ANTH and ENTH adaptors are typically generated with fluorescent proteins attached at the C-terminus, to avoid a possible obstruction of the N-terminal ANTH/ENTH domain (Song et al., 2012, Adamowski et al., 2018, Heinze et al., 2020). Yet, FP-CAP1, while likely not functional, provided surprising insights into the molecular composition of the CAP1 protein. In contrast with CAP1-mCherry, we found that FP-CAP1 expressed in *35S_pro_:GFP-CAP1* and *35S_pro_:RFP-CAP1* lines both localized to intracellular compartments, and not to the PM (Figure 3D and 3E). RFP-CAP1 co-localized at these compartments with CLC2-GFP, indicating that the intracellular binding sites of FP-CAP1 are the clathrin-positive TGNs. Surprisingly, this binding was mediated by the unstructured region of CAP1, and not by the ANTH domain, as GFP-CAP1^ΔANTH^ exhibited punctate binding patterns within the cell similar to full length GFP-CAP1, while GFP-CAP1^ΔCTERM^ was distributed in the cytosol without an association to any membranes (Figure 3E). More interestingly still, a full length GFP-CAP1 without the DPF motif (GFP-CAP1^ΔDPF^) localized at the PM in 30 of 32 seedlings, in some cases additionally associating with intracellular structures (Figure 3E), as such, presenting a localization pattern very similar to CAP1-mCherry, but not to FP-CAP1.

These observations can be interpreted as follows. FP-CAP1, in which the function of the ANTH domain is likely obstructed, preferentially associates with a clathrin-containing domain of the TGN through the unstructured region. This association is determined to a large extent by the short peptide motif DPF present in this region. In other proteins, such motifs interact with an appendage domain of AP-2 (Smith et al., 2017). Thus, it may be speculated that the binding of FP-CAP1 to the TGN is mediated by an interaction with similar appendage domains of AP-1 or AP-4, homologues of AP-2 at the TGN (Park et al., 2013; Fuji et al., 2016; Heinze et al., 2020). Furthermore, despite the presumed obstruction of the ANTH domain, FP- CAP1 can be recruited to the PM, but only when the apparently preferential binding to the TGN through the DPF motif is abolished. As such, the PM recruitment of FP- CAP1 is clearly DPF-independent, like the recruitment of CAP1-mCherry. This PM recruitment is probably not mediated by the ANTH domain, as this domain is cytosolic when expressed alone.

While these observations are difficult to translate into the native molecular function of CAP1 or ECA4, they provide evidence for the functionality of the DPF motif in these proteins. Additionally, they indicate an interesting possibility that the TGN-localized AP-1 or AP-4 adaptor complexes might engage in interactions with DPF motifs, similarly to AP-2 in non-plant systems.

### No apparent function of a double serine phosphorylation site in CAP1 and ECA4

A recent phosphoproteomic analysis of the early responses of *A. thaliana* seedling roots to treatments with auxin (Han et al., 2021) detected an auxin-induced phosphorylation at ^205^Ser in CAP1. This residue is part of a double serine phosphorylation site ^205^SRS, conserved in ECA4 as ^219^SRS (Suppl. Figure 1), and identified in a number of other proteomic studies, as well (Heazlewood et al., 2008; Curran et al., 2011; Willems et al., 2019). To assess a potential function of this phosphorylation site in the activity of CAP1 at the PM, we generated phosphorylation-mimicking and phosphorylation-deficient mutants of CAP1-mCherry.

The two serines were substituted by aspartic acid or glutamic acid (CAP1^Asp^ and CAP1^Glu^), whose negatively charged carboxyl groups imitate the negative charge of the phosphate group, as well as by alanines (CAP1^Ala^), in which case, the potential for phosphorylation is abolished. Phosphorylation mutants were expressed in *CAP1_pro_:CAP1^Asp^-mCherry, CAP1_pro_:CAP1^Glu^-mCherry,* and *CAP1_pro_:CAP1^Ala^- mCherry* lines in *cap1-1 eca4-2* knockout mutant background (described below). TIRF imaging of these CAP1-mCherry variants did not reveal any differences in their recruitment to CCPs at the PM compared to the wild type variant, as all three phosphorylation mutants were observed as distinct spots of density and persistence at the PM similar to the control (Figure 4A and 4B). As with the wild type CAP1- mCherry (Figure 1C), both phosphorylation-mimicking and phosphorylation-deficient variants could be occasionally seen exhibiting initial bursts of high recruitment to the PM followed by a decrease of signal level (Figure 4B, insets), but in most cases this pattern was not observed. Taken together, the analysis of the ^205/219^SRS phosphorylation site in CAP1 and ECA4 did not indicate its involvement in the regulation of the adaptor activity at PM-localized CCPs.

**Figure 4.**
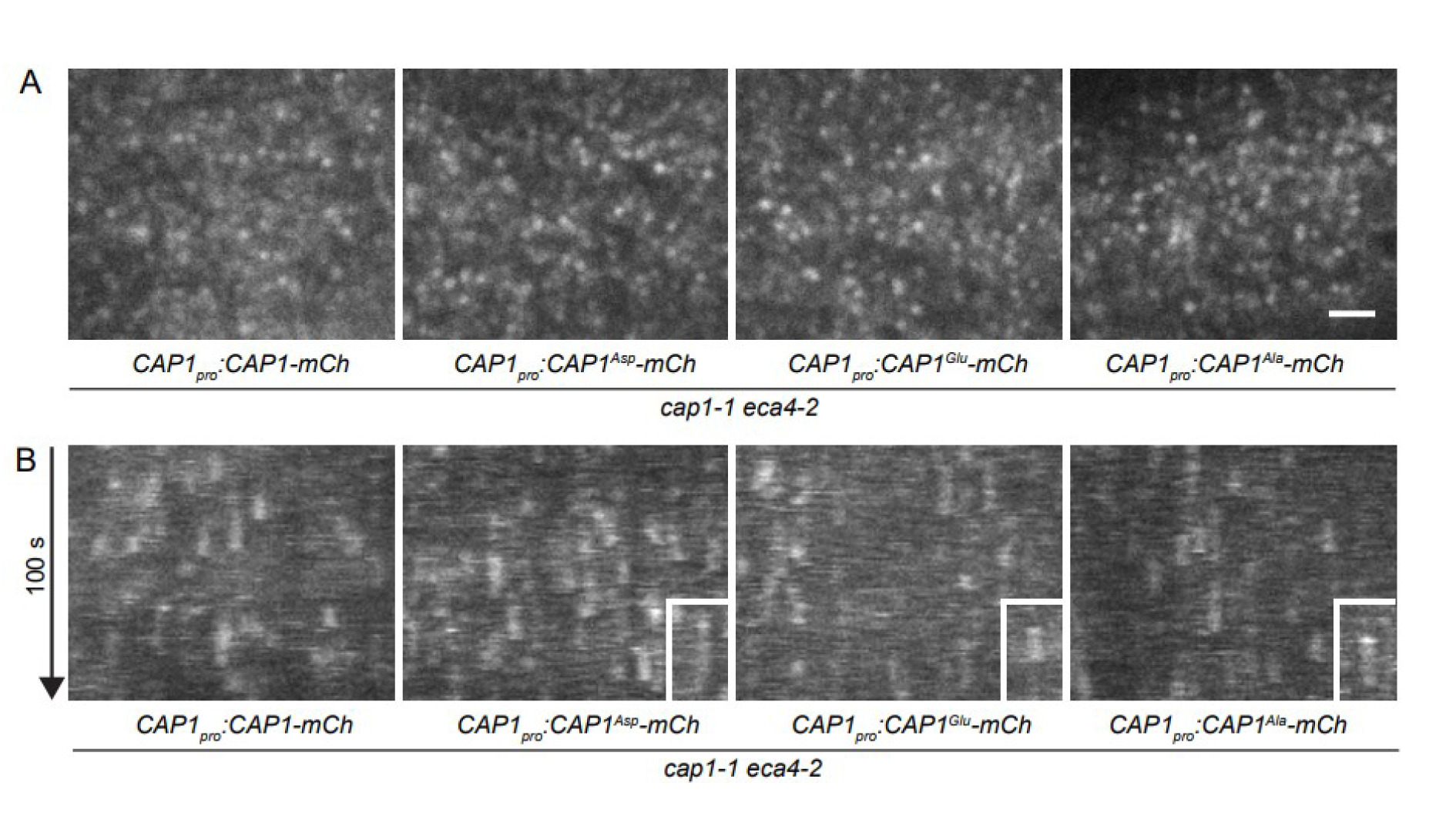
No detectable influence of a double serine phosphorylation site in CAP1 on its localization and dynamics at the PM. TIRF images (A) and kymographs of TIRF time lapses (B) of CAP1-mCherry and phosphorylation mutants CAP1^Asp^-mCherry, CAP1^Glu^-mCherry, and CAP1^Ala^-mCherry expressed in *cap1-1 eca4-2* mutant background, taken in etiolated hypocotyl epidermis. Mutations mimicking constitutive phosphorylation or abolishing phosphorylation do not influence CAP1 recruitment to the PM. Insets in (B) show rarely observed events of early bursts of CAP1 phospho-mutant recruitment to endocytic events. Scale bar – 2 µm.

### Isolation and characterization of *cap1 eca4* double mutants

On the morphological level, the previously isolated *eca4* mutants are characterized by an increase in root lengths and by a reduced sensitivity to osmotic stress (Nguyen et al., 2018). On the cellular level, they may be defective in trafficking from the TGN to the PM, as indicated by experiments involving brefeldin A, an ARF-GEF (ARF-Guanine nucleotide Exchange Factor) inhibitor, and by mislocalization of the abscisic acid transporter ABCG25 (Nguyen et al., 2018). We aimed at testing the requirement of CAP1 and ECA4 for CME by the isolation of *cap1 eca4* double mutants and by subsequent imaging of fluorescent protein fusions of molecular players involved in endocytosis using TIRF microscopy. With CRISPR/Cas9-mediated mutagenesis, we generated a *cap1* knockout allele, *cap1-1*, harboring a single nucleotide insertion shortly after the start codon of the *CAP1* gene. We crossed *cap1-1* with the two previously described T-DNA insertional mutants of *eca4*, *ateca4-1* and *ateca4-2* (Nguyen et al., 2018; herein *eca4-1* and *eca4-2*). The double mutants developed normally both at seedling and at adult stages (Figure 5A, 5B), like the single mutants (Suppl. Figure 2A, 2B), not indicative of a major developmental function of the two adaptors.

**Figure 5.**
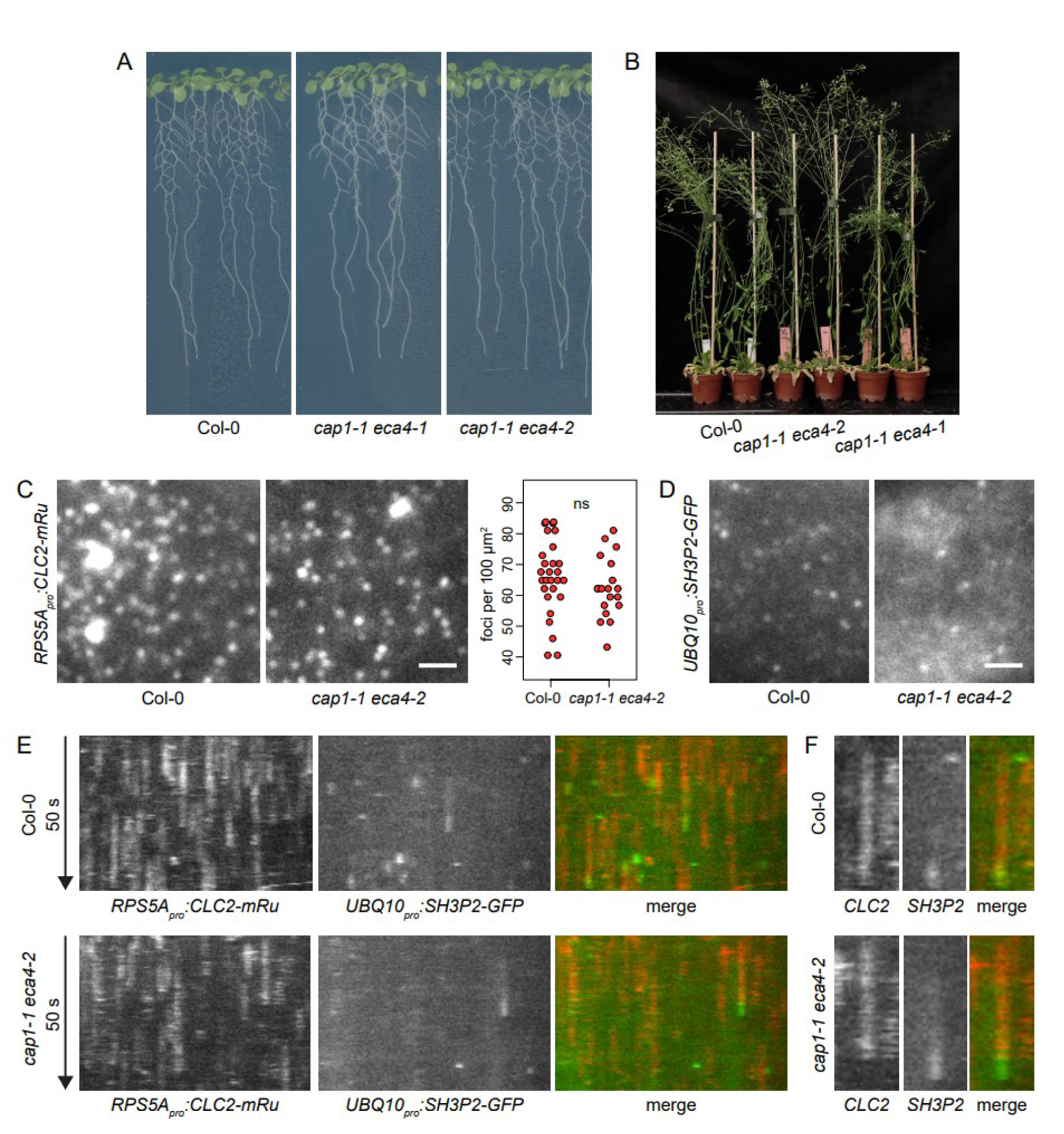
Normal development and endocytosis in *cap1 eca4* mutants. (A) Seedling development of *cap1 eca4* double mutants. (B) Adult development of *cap1 eca4* double mutants. (C) TIRF images of CLC2-mRuby3 in etiolated hypocotyl epidermis of wild type and *cap1-1 eca4-2* mutants. Scale bar – 2 µm. Graph shows a quantification of CLC2-mRuby3 foci density, each data point represents one cell. Col-0: 24.6 ± 4.7 foci per 100 µm^2^, n=28; *cap1-1 eca4-2:* 23.1 ± 3.6 foci per 100 µm^2^, n=19. Values were compared using a *t* test, ns – not significant. (D) TIRF images of SH3P2-GFP in etiolated hypocotyl epidermis of wild type and *cap1-1 eca4-2* mutants showing normal recruitment of the marker to the PM. Scale bar – 2 µm. (E, F) Kymographs of TIRF time lapse co-localizations of CLC2-mRuby3 and SH3P2-GFP in etiolated hypocotyl epidermis of wild type and *cap1-1 eca4-2* mutants. Dynamics of the PM recruitment of CLC2-mRuby3 and SH3P2-GFP are normal. (F) Detail of rarely observed events of SH3P2 recruitment at the end of CCP formation in the wild type and in *cap1-1 eca4-2*.

To assess the requirement of CAP1 and ECA4 in the endocytic process, we crossed *cap1-1 eca4-2* with a double fluorescent reporter line, *UBQ10_pro_:SH3P2- GFP RPS5A_pro_:CLC2-mRuby3* (Adamowski et al., 2022). CLC2-mRuby3, a clathrin reporter, was used to visualize the rates at which CCPs are formed at the PM in the mutant. Using TIRF microscopy, we found that the densities of CCPs at the PMs of *cap1-1 eca4-2* cells were normal, both in the epidermis of etiolated hypocotyls (Figure 5C), and in the epidermis of the elongation zone of the root (Figure 6C). The temporal development of individual CCPs in the hypocotyl appeared normal as well (Figure 5E). As such, we found no evidence for the requirement for CAP1 and ECA4 in the normal function of CME.

**Figure 6.**
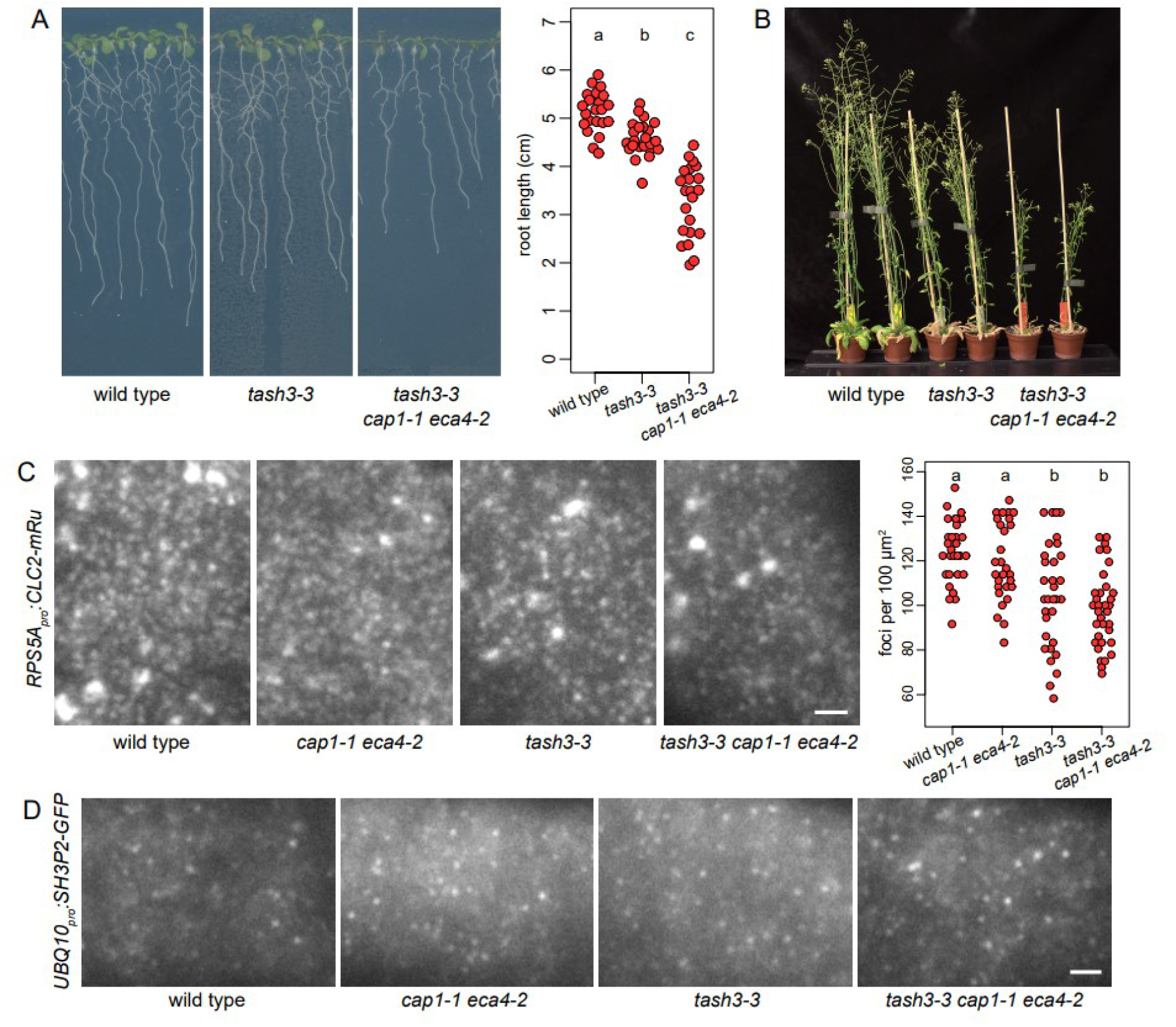
Genetic interactions between *cap1 eca4* and *tash3-3*. (A) Seedling development of *tash3-3 cap1-1 eca4-2* mutants. Lines are in crosses with *UBQ10pro:SH3P2-GFP RPS5Apro:CLC2-mRuby3*. Graph shows measurements of main root lengths of individual seedlings after 9 d of *in vitro* growth from a representative experiment. Wild type: 5.15 ± 0.41 cm, n=23; *tash3-3:* 4.58 ± 0.36 cm, n=23; *tash3-3 cap1-1 eca4-2:* 3.28 ± 0.74 cm, n=22. Values were compared using One-way ANOVA (P<0.00001) with post-hoc Tukey HSD test, groups of significantly different values are indicated. (B) Adult development of *tash3-3 cap1-1 eca4-2* mutants. Lines are in crosses with *UBQ10pro:SH3P2- GFP RPS5Apro:CLC2-mRuby3*. The triple mutant exhibited decreased growth in comparison with *tash3-3*. (C) TIRF images of CLC2-mRuby3 in the epidermis of early elongation zone of seedling roots of *tash3-3 cap1-1 eca4-2* and lower order mutants. Scale bar – 2 µm. Graph shows a quantification of CLC2-mRuby3 foci density, each data point represents one root. Wild type: 124.4 ± 13.8 foci per 100 µm^2^, n=30; *cap1-1 eca4-2:* 119.4 ± 17.4 foci per 100 µm^2^, n=28; *tash3-3:* 104.9 ± 23.2 foci per 100 µm^2^, n=32; *tash3-3 cap1-1 eca4-2:* 98.3 ± 17.0 foci per 100 µm^2^, n=34. Values were compared using One-way ANOVA (P<0.00001) with post-hoc Tukey HSD test, groups of significantly different values are indicated. (D) TIRF images of SH3P2-GFP in the epidermis of early elongation zone of seedling roots of *tash3-3 cap1-1 eca4-2* and lower order mutants. Scale bar – 2 µm.

SH3P2 is a BAR and SH3 domain protein which, together with its homologues SH3P1 and SH3P3, has a function in CME including, but not limited to, the PM recruitment of the putative CCV uncoating factors AUXILIN-LIKE1/2 (Adamowski et al., 2022). Considering that ANTH and ENTH proteins in non-plant systems contribute to membrane curvature generation during the formation of CCPs (Ford et al., 2002; Miller et al., 2015), and that SH3P1-3, like their non-plant counterparts, likely bind to the PM through curvature-sensitive BAR domains (Itoh and De Camilli, 2006; Simunovic et al., 2015; Nagel et al., 2017), the SH3P2-GFP marker was introduced into *cap1-1 eca4-2* to test the possibility that curvature-generating properties of CAP1 and ECA4 may be required for an efficient recruitment of SH3Ps to the PM. However, in *cap1-1 eca4-2*, SH3P2-GFP exhibited a normal recruitment pattern to the PM, both in the epidermis of etiolated hypocotyls (Figure 5D) and in roots (Figure 6D). Similarly to the wild type, rare events of a recruitment of SH3P2- GFP to CCPs at the conclusion of vesicle formation could be observed in *cap1-1 eca4-2* (Figure 5E and 5F; Adamowski et al., 2022).

Taken together, no evident function of CAP1 and ECA4 in the endocytic process can be detected based on the normal development, and the normal endocytosis, in *cap1 eca4* mutant plants.

**Suppl. Figure 2.**
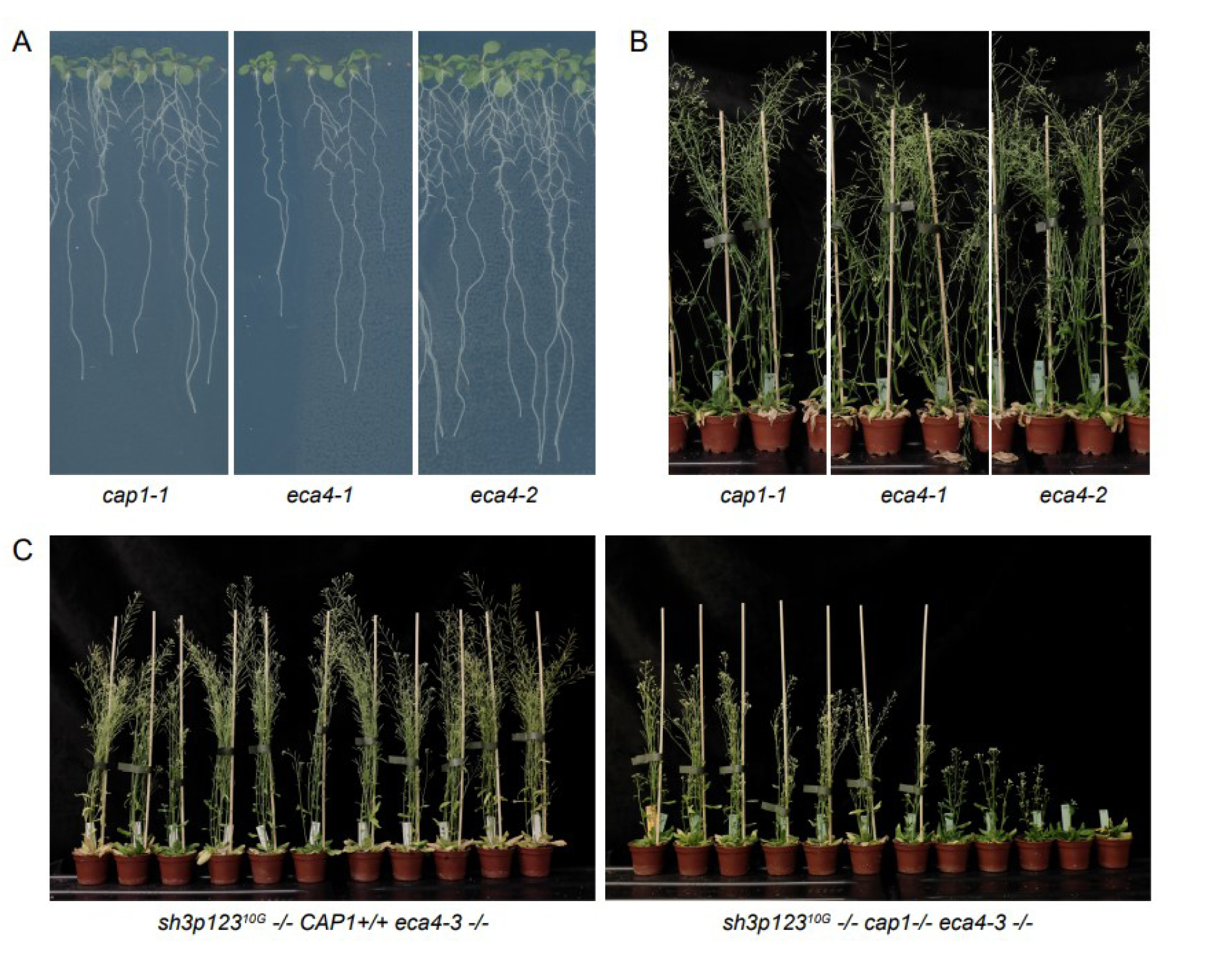
Additional mutant data. (A) Seedling development of *cap1* and *eca4* single mutants. (B) Adult development of *cap1* and *eca4* single mutants. (C) Comparison of adult development of *sh3p123^10G^ eca4-3* and *sh3p123^10G^ cap1 eca4-3.* In the right panel, plants carrying two CRISPR-induced *cap1* knockout alleles which segregated in the population are shown together, and plants are arranged according to phenotypic intensity.

### A function of CAP1 and ECA4 in CME revealed in sensitized mutant backgrounds

Mutants lacking the BAR-SH3 domain proteins SH3P1-3 exhibit a decrease in overall growth and fertility as well as in the recruitment of AUXILIN-LIKE1 to the PM, but do not exhibit measurable deficiencies in overall CME (Adamowski et al., 2022). Similarly, *tash3-3,* a partial and viable loss-of-function allele of the TPLATE complex subunit TASH3 (TPLATE-associated SH3 domain containing protein; Gadeyne et al., 2014) is characterized by a moderate reduction in growth and fertility (Adamowski et al., 2022). In turn, *sh3p123 tash3-3* quadruple homozygotes exhibit a very strong defect in overall growth, are infertile, and exhibit clearly decreased rates of CCP formation (Adamowski et al., 2022). This genetic interaction offers evidence for a function of SH3P1-3 in CME, which manifests clearly in a sensitized mutant background.

Following this example, we introduced *cap1* and *eca4* mutations into *tash3-3* and *sh3p123* mutants, expecting that a potential contribution of CAP1 and ECA4 to the endocytic process might become revealed.. By crossing, we generated a *tash3-3 cap1-1 eca4-2* triple mutant carrying *UBQ10_pro_:SH3P2-GFP* and *RPS5A_pro_:CLC2- mRuby3* reporters. Both at seedling and adult stages of development, the triple mutant exhibited reduced growth when compared with *tash3-3* (Figure 6A, 6B), as such revealing a quantitative contribution of CAP1 and ECA4 to the processes mediated by TASH3, and by extension, by the TPLATE complex, most likely to endocytosis. However, this was not detectable by TIRF microscopy of CME in the seedling root, as imaging of CLC2-mRuby3 did not reveal any changes in the moderately decreased densities of CCPs characteristic for *tash3-3* alone (Figure 6C). Similarly, the distribution of SH3P2-GFP on the PMs appeared normal both in *tash3-3* and in *tash3-3 cap1-1 eca4-2* (Figure 6D).

To combine *cap1* and *eca4* mutations with *sh3p123,* we performed *de-novo* CRISPR/Cas9 mutagenesis, rather than crossing, not only due to the complex architecture of the mutant, but also due to a close physical distance between *CAP1* and *SH3P2* loci on chromosome 4 (At4g32285 and At4g34660, respectively). Insertions of 2 and 1 nucleotides early in the coding sequences of *CAP1* and *ECA4,* respectively, were produced in the *sh3p123^10G^ CLC2_pro_:CLC2-GFP UBQ10_pro_:mCherry-AUXILIN-LIKE1* background (Adamowski et al., 2022). The resulting *sh3p123^10G^ cap1-2 eca4-3* quintuple mutant was characterized by a reduction in seedling growth which exceeded that of *sh3p123^10G^* (Figure 7A). The mutant also exhibited a variable deficiency in adult development and fertility, also exceeding in severity the phenotype of *sh3p123^10G^* (Figure 7B). This was strictly a result of combined mutation of *cap1* and *eca4,* since the phenotype was absent in *sh3p123^10G^ eca4-3*, as observed in a population segregating for *cap1* during selection (Suppl. Figure 2C). TIRF imaging of *UBQ10_pro_:mCherry-AUXILIN-LIKE1* was not conducted due to excessive silencing, but imaging of *CLC2_pro_:CLC2-GFP* in the early elongation zone of seedling roots revealed a decrease in the density of CCPs forming at the PM in *sh3p123^10G^ cap1-2 eca4-3*, contrasted with normal CCP density in *sh3p123^10G^* (Figure 7C). This result provides evidence for a role of CAP1 and ECA4 adaptors in CME, whose contribution revealed itself in an otherwise partially deficient *sh3p123* mutant background.

**Figure 7.**
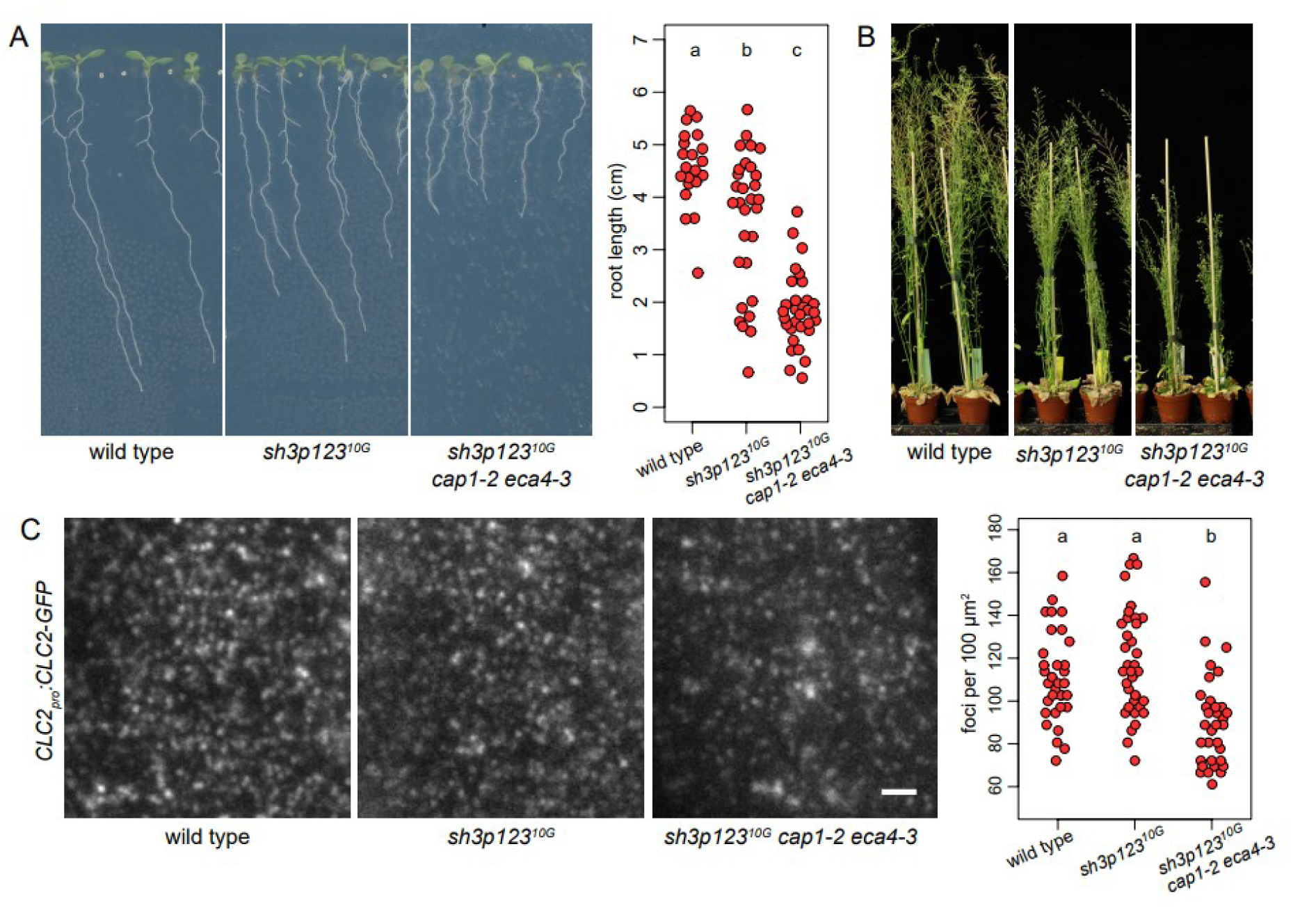
Genetic interactions between *cap1 eca4* and *sh3p123*. (A) Seedling development of *sh3p123^10G^ cap1-2 eca4-3* and *sh3p123^10G^*. All lines contain *CLC2pro:CLC2-GFP UBQ10pro:mCherry-AUXILIN-LIKE1*. Graph shows measurements of main root lengths of individual seedlings after 9 d of *in vitro* growth from a representative experiment. Wild type: 4.57 ± 0.73 cm, n=21; *sh3p123^10G^:* 3.57 ± 1.31 cm, n=30; *sh3p123^10G^ cap1-2 eca4-3:* 1.85 ± 0.69 cm, n=32. Values were compared using One-way ANOVA (P<0.00001) with post-hoc Tukey HSD test, groups of significantly different values are indicated. (B) Adult development of *sh3p123^10G^ cap1-2 eca4-3* mutant. All lines contain *CLC2pro:CLC2-GFP UBQ10pro:mCherry-AUXILIN-LIKE1*. The quintuple mutant has a variable deficiency in overall growth exceeding in degree that of *sh3p123^10G^*. (C) TIRF images of CLC2-GFP in the epidermis of early elongation zone of seedling roots of *sh3p123^10G^ cap1-2 eca4-3* and *sh3p123^10G^*. Scale bar – 2 µm. Graph shows a quantification of CLC2- GFP foci density, each data point represents one root. Wild type: 111.1 ± 20.5 foci per 100 µm^2^, n=33; *sh3p123^10G^:* 118.3 ± 24.4 foci per 100 µm^2^, n=35; *sh3p123^10G^ cap1-2 eca4-3:* 90.2 ± 20.8 foci per 100 µm^2^, n=33. Values were compared using One-way ANOVA (P<0.00001) with post-hoc Tukey HSD test, groups of significantly different values are indicated.

In summary, by introducing *cap1* and *eca4* mutations into sensitized *tash3-3* and *sh3p123* mutant backgrounds, we were able to reveal a contribution of CAP1 and ECA4 in CME, manifested by aggravated developmental deficiencies of the resulting mutants, and, in the case of *sh3p123*, also in an aggravation of deficiencies in CME measured by fluorescent microscopy.

## Discussion

Adaptor proteins serve key functions during the formation of clathrin-coated vesicles (Owen et al., 2004). The ANTH/ENTH/VHS superfamily of monomeric adaptor proteins is expanded in plants (Zouhar and Sauer, 2014), and the characterization of *A. thaliana* homologues has been a matter of increasing interest in the recent years (Muro et al., 2018; Heinze et al., 2020; Fujimoto et al., 2020). Here, we contribute to this effort with a molecular and functional characterization of the homologous CAP1 and ECA4 ANTH adaptor pair.

High sequence similarity suggests a redundant function of CAP1 and ECA4 (Suppl. Figure 1). This similarity extends to the unstructured tail of the two adaptors, whose length and composition is otherwise variable between monomeric adaptor isoforms (Zouhar and Sauer, 2014). We tested the function of a double serine phosphorylation site in this region of both adaptors, but its mutation in CAP1- mCherry did not influence the recruitment dynamics of the adaptor to the PM. In turn, we identified a conserved DPF motif, which, while not needed for adaptor recruitment to the PM, manifested its functional importance by affecting the localization of an unusual, N-terminal fluorescent protein fusion of CAP1.

ECA4-GFP localizes both to the PM and to the TGN, and ECA4 has been previously ascribed a function in trafficking from the latter compartment, based on single mutant study (Nguyen et al., 2018). In turn, despite high sequence similarity, CAP1-mCherry persistently localizes to the PM, while it is observed at intracellular structures only rarely (Adamowski et al., 2018). It is an open question whether localization differences of the fluorescent reporters reflect true differences in the sites of action of the native proteins, but it may be, indeed, that the two proteins are not identical entirely in molecular functions. The analysis of *cap1-1* as well as *cap1 eca4* in the context of TGN-to-PM trafficking would be informative in this context. In this connection, we note the existence of glycine rich insertions in the unstructured tail of ECA4 (Suppl. Figure 1), which may alter the molecular interactions of ECA4 in comparison with CAP1.

Regardless of these possible differences, we identify here a redundant function of the two adaptors in CME. The specific pattern of CAP1 recruitment to CCPs at the PM, characterized by a strong initial burst of binding during early steps of clathrin polymerization, would represent, to our knowledge, a novel temporal pattern of endocytic protein recruitment, not found in studies in plant and non-plant systems (Taylor et al, 2011). However, we treat this observation with reserve due to its rarity when CAP1 or ECA4 fluorescent markers were expressed without CLC2-GFP. The reason for the variation in the observations are not clear. Regardless of this, the increased recruitment of CAP1-mCherry to the PM following overexpression of AUXILIN-LIKE1, similar to the behavior of subunits of the early-arriving TPLATE and AP-2 complexes (Adamowski et al., 2018; Gadeyne et al., 2014), supports the notion that CAP1 as well as ECA4, participate in vesicle formation from its early stages.

Overall, our study indicates only a minor contribution of CAP1 and ECA4 adaptors to the endocytic process, at least in optimal growth conditions. Double knockout mutants develop correctly, and do not manifest deficiencies in CME in seedlings, but deficiencies resulting from the absence of these proteins are manifested in mutant backgrounds sensitized in terms of the endocytic process. It is possible that other ANTH family adaptors, less conserves in sequence, share a redundant function with CAP1 and ECA4, although the specific identification of CAP1 and/or ECA4, but not other monomeric adaptors, in interaction studies of the TPLATE complex (Gadeyne et al., 2014), and of CLC (Adamowski et al., 2018), is indicative of a unique function of CAP1 and ECA4, reflected in this case by unique molecular interactions. It is rather more likely that these ANTH adaptors simply play only a minor, accessory role in the formation of endocytic CCVs.

The formation of CCVs should be, perhaps, perceived not as analogical to a linear signaling pathway, with a number of essential factors acting in succession, but more as a collection of distinct proteins which, in a facultative, or merely quantitative sense, contribute to the shaping of the vesicle. In this scenario, various proteins are able to support the successful outcome through mechanistically distinct activities. The observation that developmental and cellular phenotypes manifest in mutants of two distinct endocytic protein classes, but are absent or poorly manifested when any one class is missing (Adamowski et al., 2022 and this study), supports this perception.

## Materials and Methods

### Plant material

The following previously described *A. thaliana* lines were used in this study: *CsVMV_pro_:ECA4-GFP* (Song et al., 2012), *eca4-1* (SALK_114631C; Nguyen et al., 2018)*, eca4-2* (GK_839F07; Nguyen et al., 2018), *CLC2_pro_:CLC2-GFP* (Konopka et al., 2008), *UBQ10_pro_:SH3P2-GFP RPS5A_pro_:CLC2-mRuby3*, *tash3-3*, *sh3p1,2,3^10G-4E^ CLC2_pro_:CLC2-GFP UBQ10_pro_:mCh-AUXILIN-LIKE1* (Adamowski et al., 2022), *CAP1_pro_:CAP1-mCherry*, *35S_pro_:CAP1-RFP*, *XVE»AUXILIN-LIKE1* (Adamowski et al., 2018), Lines generated as part of this study are listed in Suppl. Table 1 and primers used for genotyping in Suppl. Table 2.

### *In vitro* cultures of *Arabidopsis* seedlings

Seedlings were grown in *in vitro* cultures on half-strength Murashige and Skoog (½MS) medium of pH=5.9 supplemented with 1% (w/v) sucrose and 0.8% (w/v) phytoagar at 21°C in 16h light/8h dark cycles with Philips GreenPower LED as light source, using deep red (660nm)/far red (720nm)/blue (455nm) combination, with a photon density of about 140µmol/(m^2^s) +/- 20%. β-estradiol (Sigma-Aldrich) was solubilized in 100% ethanol to 5 mg/mL stock concentration and added to ½MS media during preparation of solid media to a final concentration of 2.5 µg/mL.

Seedlings of *XVE»AUXILIN-LIKE1* lines were induced by transferring to β-estradiol-supplemented media at day 3. Petri dishes for TIRF imaging in hypocotyls of etiolated seedlings were initially exposed to light for several hours and then wrapped in aluminium foil.

### Confocal Laser Scanning Microscopy

4 to 5 d old seedlings were used for live imaging with Zeiss LSM800 confocal laser scanning microscope with 20X lens. Measurements of PM signal intensity of CAP1- mCherry was performed in Fiji (https://imagej.net/Fiji) as a mean grey value of a line of 5 pixel width drawn over multiple PMs in each CLSM image.

### Total Internal Reflection Fluorescence microscopy

Early elongation zone of roots in excised ∼1 cm long root tip fragments from 7d old seedlings, as well as apical ends of excised hypocotyls from 3d old etiolated seedlings, were used for TIRF imaging. Imaging was performed with Olympus IX83 TIRF microscope, using a 100X TIRF lens with an additional 1.6X or 2X magnification lens in the optical path. Time lapses of 100 frames at 0.5 s or 1 s intervals with exposure times of 195 ms or 200 ms, or single snapshots of 200 ms exposure, were taken, depending on the experiment. Two-channel time lapses were captured sequentially. CLC2-GFP and CLC2-mRuby3 foci were counted in square regions of 36 µm^2^ taken from the captured TIRF images or movies using Fiji (https://imagej.net/Fiji).

### Molecular cloning

All constructs generated in this study are listed in Suppl. Table 3 and primers used for cloning in Suppl. Table 2. CAP1 truncations in variants with and without stop codons were cloned into pDONR221 Gateway entry vectors (Invitrogen) by BP Clonase. DPF motif and phosphorylation site mutagenesis in CAP1 were performed by overlap-extension PCR with mutations introduced on overlapping internal primers. ^555^DPF was mutated to ^555^AAA by substituting ^1663^GACCCGTTC for ^1663^GCCGCGGCC. The double serine phosphorylation site ^205^SRS was mutated to ^205^DRD, ^205^ERE, and ^205^ARA, by substituting ^613^TCGAGGTCT to ^613^GACAGGGAT, ^613^GAGAGGGAG, and ^613^GCGAGGGCT, respectively. Resulting mutant variants of CAP1 were cloned in versions with and without stop codons into pDONR221 Gateway entry vectors (Invitrogen) by BP Clonase. N-terminal GFP fusions of CAP1 variants were produced by introducing CAP1 variants from pDONR221 or pENTR/D-TOPO entry vectors into pB7WGF2 expression vectors (Karimi et al., 2002) by LR Clonase II (Invitrogen). Previously described CAP1/pENTR/D-TOPO (Adamowski et al., 2018) was used for wild type constructs. 35S:RFP-CAP1 was produced similarly by introducing CAP1 from CAP1/pENTR/D-TOPO into pK7WGR2 (Karimi et al., 2002). C-terminal mCherry fusions of CAP1 variants were produced by combining ProCAP1/pDONRP4P1r (Adamowski et al., 2018), CAP1 mutant variants in pDONR221, and mCherry/pDONRP2rP3 into pK7m34GW (phospho-mutants) or pH7m34GW (truncations and DPF deletion) expression vectors (Karimi et al., 2002) by LR Clonase II (Invitrogen).

### CRISPR/Cas9 mutagenesis

Details of generated mutant alleles are given in Suppl. Table 1. CRISPR mutagenesis of *CAP1* in Col-0 background (*cap1-1*) was performed by a previously published method (Richter et al., 2018). CRISPR mutagenesis of *CAP1* and *ECA4* in *sh3p1,2,3^10G-4E^ CLC2_pro_:CLC2-GFP UBQ10_pro_:mCh-AUXILIN-LIKE1* background was performed with the use of pHEE401 binary vector and template plasmid pCBC- DT1T2 (Wang et al., 2015). sgRNA sequences were selected with the use of CRISPR RGEN Tools website (http://www.rgenome.net/cas-designer/). The following sgRNA sequences were used: *CAP1* AAAGCGATCGGAGTTGTTA; *ECA4* ACCTGACCTCTCTCTCTCG. In T1 plants, target sequences were PCR-amplified and sequenced using primers listed in Suppl. Table 2. A single T1 plant carrying a homozygous 1xT insertion in *ECA4 (eca4-3)* and heterozygous for a 2xT insertion in *CAP1 (cap1-2)* was propagated to T2. T2 generation plants were genotyped to select *cap1-2* homozygotes. Due to a high number of inserted CRISPR/Cas9 transgenes in this line, mutant plants without the transgenes were not successfully isolated.

### Accession numbers

Sequence data from this article can be found in the GenBank/EMBL libraries under the following accession numbers: CAP1 (AT4G32285), ECA4 (AT2G25430), AUXILIN-LIKE1 (AT4G12780), TASH3 (AT2G07360), SH3P1 (AT1G31440), SH3P2 (AT4G34660), SH3P3 (AT4G18060), CLC2 (AT2G40060).

## Author Contributions

M.A. and J.F. designed research, analyzed data and wrote the manuscript. M.A. an I.M. performed research.

## Acknowledgements

The authors wish to acknowledge James Matthew Watson, Monika Borowska, and Peggy Stolt-Bergner at ProTech Facility of the Vienna Biocenter Core Facilities for CRISPR/Cas9 mutagenesis of *cap1-1,* Prof. Inhwan Hwang for sharing the *CsVMV_pro_:ECA4-GFP* reporter line, and Prof. Qi-Jun Chen for sharing plasmids used for CRISPR/Cas9 mutagenesis. This work was supported by the Austrian Science Fund (FWF): I 3630-B25.

## Supplementary Tables

**Supplementary Table 1.**
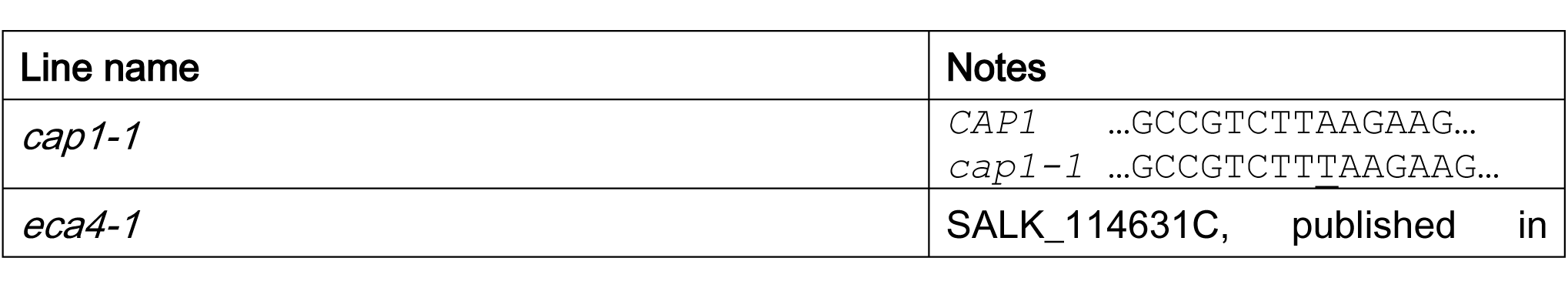
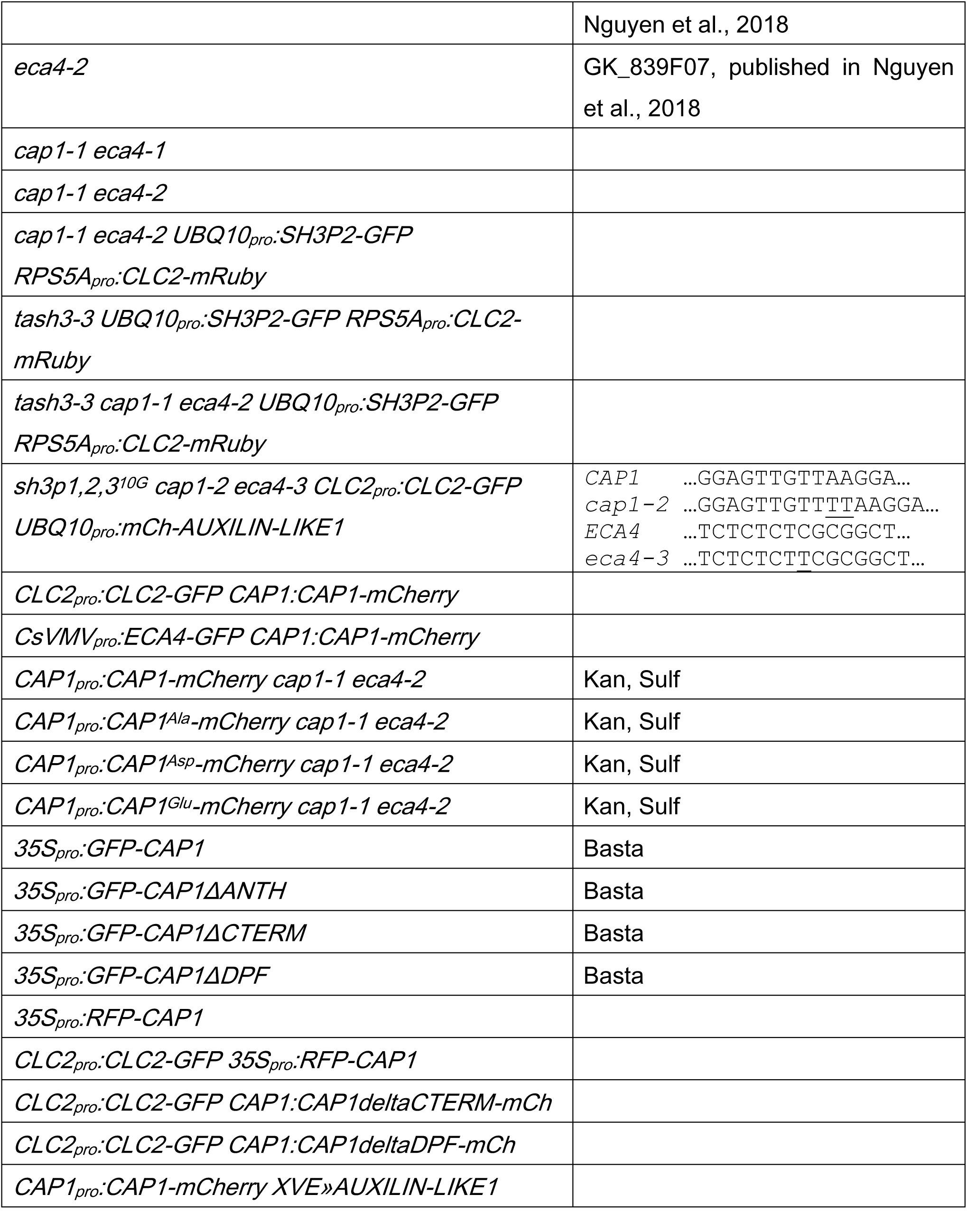
Lines generated as part of this study.

**Supplementary Table 2.**
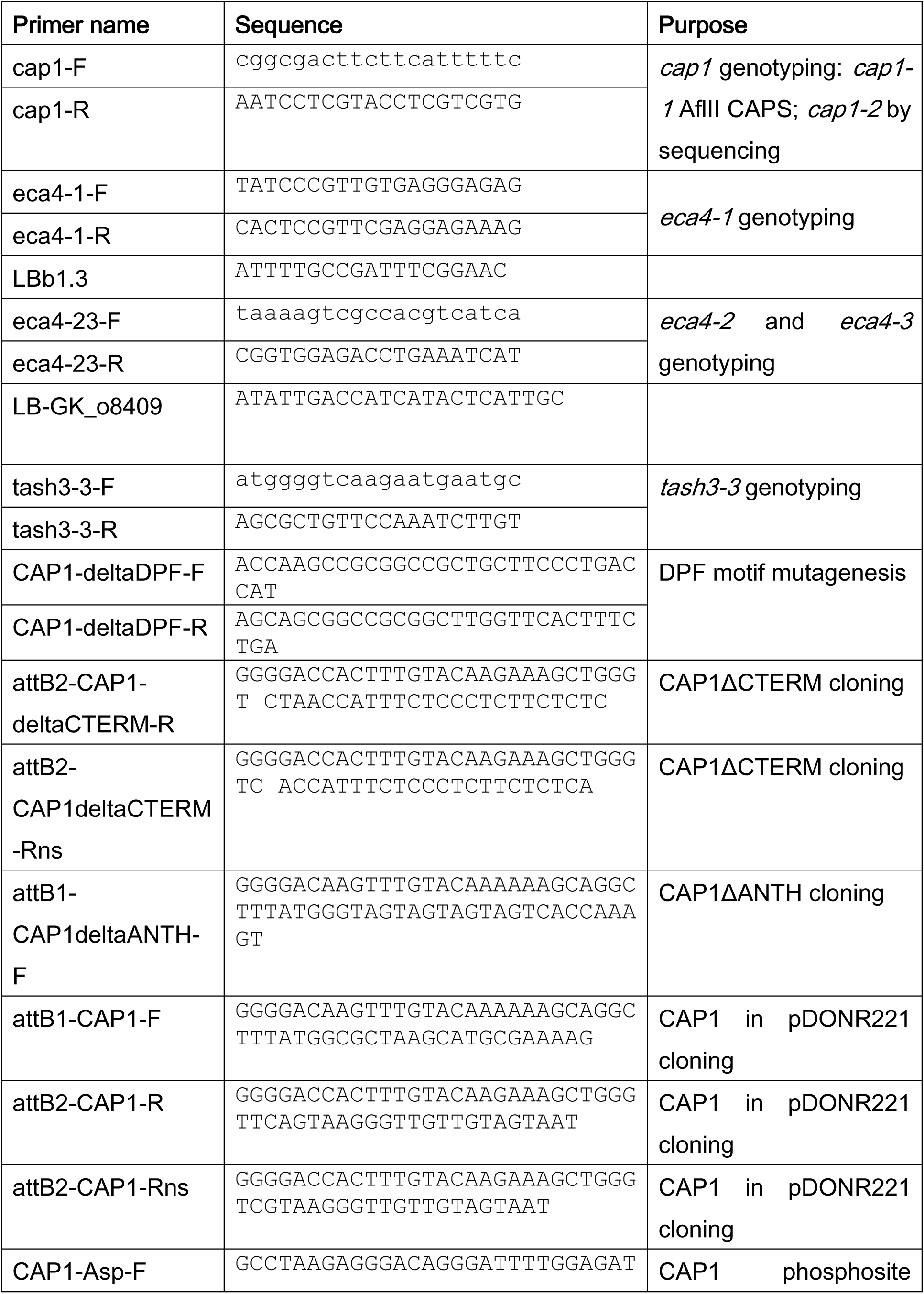
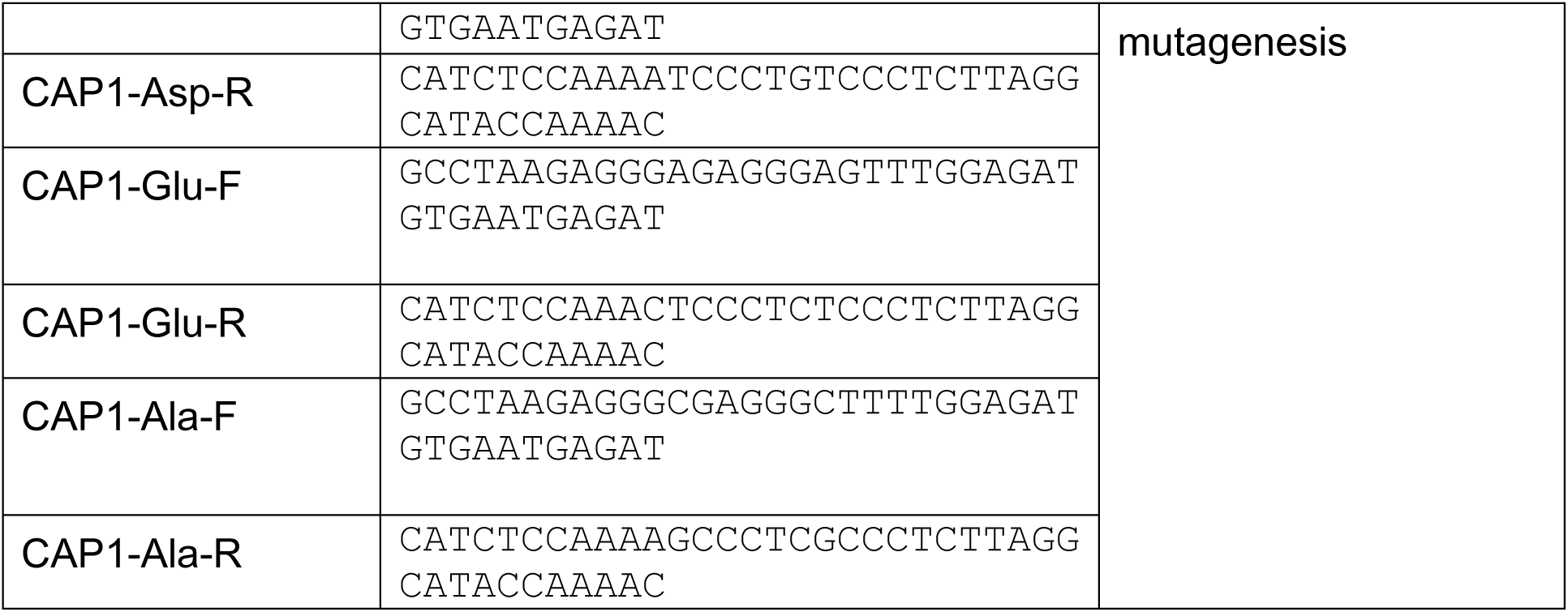
Primers used in this study.

**Supplementary Table 3.** Plasmids generated in this study.

| Plasmid name | Notes |
| --- | --- |
| CAP1_deltaANTH/pDONR221 | with STOP codon |
| CAP1_deltaANTH/pDONR221 | without STOP codon |
| CAP1_deltaCTERM/pDONR221 | with STOP codon |
| CAP1_deltaCTERM/pDONR221 | without STOP codon |
| CAP1_deltaDPF/pDONR221 | with STOP codon |
| CAP1_deltaDPF/pDONR221 | without STOP codon |
| CAP1_Ala/pDONR221 | with STOP codon |
| CAP1_Ala/pDONR221 | without STOP codon |
| CAP1_Glu/pDONR221 | with STOP codon |
| CAP1_Glu/pDONR221 | without STOP codon |
| CAP1_Asp/pDONR221 | with STOP codon |
| CAP1_Asp/pDONR221 | without STOP codon |
| 35S:RFP-CAP1/pK7WGR2 |  |
| 35S:GFP-CAP1/pB7WGF2 |  |
| 35S:GFP-CAP1_deltaANTH/pB7WGF2 |  |
| 35S:GFP-CAP1_deltaCTERM/pB7WGF2 |  |
| 35S:GFP-CAP1_deltaDPF/pB7WGF2 |  |
| CAP1:CAP1_Ala-mCh/pK7m34GW |  |
| CAP1:CAP1_Asp-mCh/pK7m34GW |  |
| CAP1:CAP1_Glu-mCh/pK7m34GW |  |
| CAP1:CAP1_deltaANTH-mCh/pH7m34GW |  |
| CAP1:CAP1_deltaCTERM-mCh/pH7m34GW |  |
| CAP1:CAP1_deltaDPF-mCh/pH7m34GW |  |
| CAP1ECA4 CRISPR/pHEE401 | Used for mutagenesis in *sh3p123* background |

## Notes

### Competing Interest Statement

The authors have declared no competing interest.

## References

Adamowski, M., Matijević, I., and Friml, J. (2021) The role of clathrin in exocytosis and the mutual regulation of endo- and exocytosis in plant cells. BIORXIV/2021/468992

Adamowski, M., Matijević, I., Narasimhan, M., and Friml, J. (2022) SH3Ps recruit auxilin-like vesicle uncoating factors into clathrin-mediated endocytosis BIORXIV/2022/475403

Adamowski, M., Narasimhan, M., Kania, U., Glanc, M., De Jaeger, G., and Friml, J. (2018). A Functional Study of AUXILIN-LIKE1 and 2, Two Putative Clathrin Uncoating Factors in Arabidopsis. Plant Cell 30: 700–716.

Béthune, J. and Wieland, F.T. (2018). Assembly of COPI and COPII Vesicular Coat Proteins on Membranes. Annu. Rev. Biophys. 47: 63–83.

Busch, D.J., Houser, J.R., Hayden, C.C., Sherman, M.B., Lafer, E.M., and Stachowiak, J.C. (2015). Intrinsically disordered proteins drive membrane curvature. Nat Commun 6: 7875.

Curran, A., Chang, I.-F., Chang, C.-L., Garg, S., Miguel, R.M., Barron, Y.D., Li, Y., Romanowsky, S., Cushman, J.C., Gribskov, M., Harmon, A.C., and Harper, J.F. (2011) Calcium-Dependent Protein Kinases from Arabidopsis Show Substrate Specificity Differences in an Analysis of 103 Substrates. Front. Plant Sci. 2.

Ford, M.G.J., Mills, I.G., Peter, B.J., Vallis, Y., Praefcke, G.J.K., Evans, P.R., and McMahon, H.T. (2002). Curvature of clathrin-coated pits driven by epsin. Nature 419: 361–366.

Fotin, A., Cheng, Y., Sliz, P., Grigorieff, N., Harrison, S.C., Kirchhausen, T., and Walz, T. (2004). Molecular model for a complete clathrin lattice from electron cryomicroscopy. Nature 432: 573–579.

Fuji, K., Shirakawa, M., Shimono, Y., Kunieda, T., Fukao, Y., Koumoto, Y., Takahashi, H., Hara-Nishimura, I., and Shimada, T. (2016). The Adaptor Complex AP-4 Regulates Vacuolar Protein Sorting at the trans-Golgi Network by Interacting with VACUOLAR SORTING RECEPTOR1. Plant Physiol. 170: 211–219.

Fujimoto, M., Ebine, K., Nishimura, K., Tsutsumi, N., and Ueda, T. (2020). Longin R-SNARE is retrieved from the plasma membrane by ANTH domain-containing proteins in *Arabidopsis*. Proc Natl Acad Sci USA 117: 25150–25158.

Gadeyne, A. et al. (2014). The TPLATE adaptor complex drives clathrin-mediated endocytosis in plants. Cell 156.

Garcia-Alai, M.M., Heidemann, J., Skruzny, M., Gieras, A., Mertens, H.D.T., Svergun, D.I., Kaksonen, M., Uetrecht, C., and Meijers, R. (2018). Epsin and Sla2 form assemblies through phospholipid interfaces. Nat Commun 9: 328.

H. Han, I. Verstraeten, M. Roosjen, E. Mazur, N. Rýdza, J. Hajný, K. Ötvös, D. Weijers, J. Friml. (2021) Rapid auxin-mediated phosphorylation of Myosin regulates trafficking and polarity in Arabidopsis. bioRxiv, 439603

Heazlewood JL, Durek P, Hummel J, Selbig J, Weckwerth W, Walther D, Schulze WX (2008) PhosPhAt: A Database of phosphorylation sites in Arabidopsis thaliana and a plant specific phosphorylation site predictor. Nucleic Acids Research 36: D1015–D1021

Heinze, L., Freimuth, N., Rößling, A.-K., Hahnke, R., Riebschläger, S., Fröhlich, A., Sampathkumar, A., McFarlane, H.E., and Sauer, M. (2020). EPSIN1 and MTV1 define functionally overlapping but molecularly distinct *trans* -Golgi network subdomains in *Arabidopsis*. Proc Natl Acad Sci USA 117: 25880–25889.

Itoh, T. (2001). Role of the ENTH Domain in Phosphatidylinositol-4,5-Bisphosphate Binding and Endocytosis. Science 291: 1047–1051.

Itoh, T. and De Camilli, P. (2006). BAR, F-BAR (EFC) and ENTH/ANTH domains in the regulation of membrane–cytosol interfaces and membrane curvature. Biochimica et Biophysica Acta (BBA) - Molecular and Cell Biology of Lipids 1761: 897–912.

Kaksonen, M. and Roux, A. (2018). Mechanisms of clathrin-mediated endocytosis. Nat Rev Mol Cell Biol 19: 313–326.

Karimi, M., Inzé, D., and Depicker, A. (2002). GATEWAY^TM^ vectors for Agrobacterium-mediated plant transformation. Trends Plant Sci. 7: 193–195.

Konopka, C.A., Backues, S.K., and Bednarek, S.Y. (2008). Dynamics of Arabidopsis Dynamin-Related Protein 1C and a Clathrin Light Chain at the Plasma Membrane. Plant Cell Online 20: 1363–1380.

Li, H., Luo, N., Wang, W., Liu, Z., Chen, J., Zhao, L., Tan, L., Wang, C., Qin, Y., Li, C., Xu, T., and Yang, Z. (2018). The REN4 rheostat dynamically coordinates the apical and lateral domains of Arabidopsis pollen tubes. Nat Commun 9: 2573.

Miller, S.E., Collins, B.M., McCoy, A.J., Robinson, M.S., and Owen, D.J. (2007). A SNARE–adaptor interaction is a new mode of cargo recognition in clathrin-coated vesicles. Nature 450: 570–574.

Miller, S.E., Mathiasen, S., Bright, N.A., Pierre, F., Kelly, B.T., Kladt, N., Schauss, A., Merrifield, C.J., Stamou, D., Höning, S., and Owen, D.J. (2015). CALM Regulates Clathrin-Coated Vesicle Size and Maturation by Directly Sensing and Driving Membrane Curvature. Developmental Cell 33: 163–175.

Miller, S.E., Sahlender, D.A., Graham, S.C., Höning, S., Robinson, M.S., Peden, A.A., and Owen, D.J. (2011). The Molecular Basis for the Endocytosis of Small R-SNAREs by the Clathrin Adaptor CALM. Cell 147: 1118–1131.

Muro, K., Matsuura-Tokita, K., Tsukamoto, R., Kanaoka, M.M., Ebine, K., Higashiyama, T., Nakano, A., and Ueda, T. (2018). ANTH domain-containing proteins are required for the pollen tube plasma membrane integrity via recycling ANXUR kinases. Commun Biol 1: 152.

Narasimhan, M., Johnson, A., Prizak, R., Kaufmann, W.A., Tan, S., Casillas-Pérez, B., and Friml, J. (2020). Evolutionarily unique mechanistic framework of clathrin-mediated endocytosis in plants. eLife 9: e52067.

Nguyen, H.H., Lee, M.H., Song, K., Ahn, G., Lee, J., and Hwang, I. (2018). The A/ENTH Domain-Containing Protein AtECA4 Is an Adaptor Protein Involved in Cargo Recycling from the trans-Golgi Network/Early Endosome to the Plasma Membrane. Molecular Plant 11: 568–583.

Owen, D.J., Collins, B.M., and Evans, P.R. (2004). ADAPTORS FOR CLATHRIN COATS: Structure and Function. Annu. Rev. Cell Dev. Biol. 20: 153–191.

Park, M., Song, K., Reichardt, I., Kim, H., Mayer, U., Stierhof, Y.-D., Hwang, I., and Jurgens, G. (2013). Arabidopsis mu-adaptin subunit AP1M of adaptor protein complex 1 mediates late secretory and vacuolar traffic and is required for growth. Proc. Natl. Acad. Sci. 110: 10318–10323.

Park, S.Y. and Guo, X. (2014). Adaptor protein complexes and intracellular transport. Bioscience Reports 34: e00123.

Richter, J., Watson, J.M., Stasnik, P., Borowska, M., Neuhold, J., Berger, M., Stolt-Bergner, P., Schoft, V., and Hauser, M.-T. (2018). Multiplex mutagenesis of four clustered CrRLK1L with CRISPR/Cas9 exposes their growth regulatory roles in response to metal ions. Sci Rep 8: 12182.

Di Rubbo, S. et al. (2013). The Clathrin Adaptor Complex AP-2 Mediates Endocytosis of BRASSINOSTEROID INSENSITIVE1 in Arabidopsis. Plant Cell 25: 2986–2997.

Sanger, A., Hirst, J., Davies, A.K., and Robinson, M.S. (2019). Adaptor protein complexes and disease at a glance. J Cell Sci 132: jcs222992.

Sauer, M., Delgadillo, M.O., Zouhar, J., Reynolds, G.D., Pennington, J.G., Jiang, L., Liljegren, S.J., Stierhof, Y.-D., De Jaeger, G., Otegui, M.S., Bednarek, S.Y., and Rojo, E. (2013). MTV1 and MTV4 Encode Plant-Specific ENTH and ARF GAP Proteins That Mediate Clathrin-Dependent Trafficking of Vacuolar Cargo from the Trans-Golgi Network. The Plant Cell 25: 2217–2235.

Scheele, U., Alves, J., Frank, R., Düwel, M., Kalthoff, C., and Ungewickell, E. (2003). Molecular and Functional Characterization of Clathrin- and AP-2-binding Determinants within a Disordered Domain of Auxilin. Journal of Biological Chemistry 278: 25357–25368.

Simunovic, M., Voth, G.A., Callan-jones, A., and Bassereau, P. (2015). When Physics Takes Over: BAR Proteins and Membrane Curvature. Trends Cell Biol. 25: 780–792.

Skruzny, M. et al. (2015). An Organized Co-assembly of Clathrin Adaptors Is Essential for Endocytosis. Developmental Cell 33: 150–162.

Smith, S.M., Baker, M., Halebian, M., and Smith, C.J. (2017). Weak Molecular Interactions in Clathrin-Mediated Endocytosis. Front. Mol. Biosci. 4: 72.

Song, K., Jang, M., Kim, S.Y., Lee, G., Lee, G.-J., Kim, D.H., Lee, Y., Cho, W., and Hwang, I. (2012). An A/ENTH Domain-Containing Protein Functions as an Adaptor for Clathrin-Coated Vesicles on the Growing Cell Plate in Arabidopsis Root Cells. Plant Physiology 159: 1013–1025.

Song, J., Lee, M.H., Lee, G.-J., Yoo, C.M., and Hwang, I. (2006). *Arabidopsis* EPSIN1 Plays an Important Role in Vacuolar Trafficking of Soluble Cargo Proteins in Plant Cells via Interactions with Clathrin, AP-1, VTI11, and VSR1. Plant Cell 18: 2258–2274.

Stachowiak, J.C., Brodsky, F.M., and Miller, E.A. (2013). A cost–benefit analysis of the physical mechanisms of membrane curvature. Nat Cell Biol 15: 1019–1027.

Stachowiak, J.C., Schmid, E.M., Ryan, C.J., Ann, H.S., Sasaki, D.Y., Sherman, M.B., Geissler, P.L., Fletcher, D.A., and Hayden, C.C. (2012). Membrane bending by protein–protein crowding. Nat Cell Biol 14: 944–949.

Stahelin, R.V., Long, F., Peter, B.J., Murray, D., De Camilli, P., McMahon, H.T., and Cho, W. (2003). Contrasting Membrane Interaction Mechanisms of AP180 N-terminal Homology (ANTH) and Epsin N-terminal Homology (ENTH) Domains. Journal of Biological Chemistry 278: 28993–28999.

Stachowiak, J.C., Brodsky, F.M., and Miller, E.A. (2013). A cost–benefit analysis of the physical mechanisms of membrane curvature. Nat Cell Biol 15: 1019– 1027.

Taylor, M.J., Perrais, D., and Merrifield, C.J. (2011). A High Precision Survey of the Molecular Dynamics of Mammalian Clathrin-Mediated Endocytosis. PLoS Biol 9: e1000604.

Wang, Z.-P., Xing, H.-L., Dong, L., Zhang, H.-Y., Han, C.-Y., Wang, X.-C., and Chen, Q.-J. (2015). Egg cell-specific promoter-controlled CRISPR/Cas9 efficiently generates homozygous mutants for multiple target genes in Arabidopsis in a single generation. Genome Biol 16: 144.

Willems P, Horne A, Van Parys T, Goormachtig S, De Smet I, Botzki A, Van Breusegem F, Gevaert K. (2019) The Plant PTM Viewer, a central resource for exploring plant protein modifications. Plant J. 99(4):752–762.

Zouhar, J. and Sauer, M. (2014). Helping Hands for Budding Prospects: ENTH/ANTH/VHS Accessory Proteins in Endocytosis, Vacuolar Transport, and Secretion. Plant Cell 26: 4232–4244.

Zwiewka, M., Feraru, E., Möller, B., Hwang, I., Feraru, M.I., Kleine-Vehn, J., Weijers, D., and Friml, J. (2011). The AP-3 adaptor complex is required for vacuolar function in Arabidopsis. Cell Res. 21: 1711–1722.

